# Microglial plasticity governed by state-specific enhancer landscapes

**DOI:** 10.1101/2025.01.30.635595

**Authors:** Nicole Hamagami, Dvita Kapadia, Nora Abduljawad, Zuolin Cheng, Liam McLaughlin, Darsh Singhania, Kia M. Barclay, Jin Yang, Zhixin Sun, Peter Bayguinov, Guoqiang Yu, Harrison W. Gabel, Qingyun Li

## Abstract

Single-cell transcriptomic studies have identified distinct microglial subpopulations with shared and divergent gene signatures across development, aging and disease. Whether these microglial subsets represent ontogenically separate lineages of cells, or they are manifestations of plastic changes of microglial states downstream of some converging signals is unknown. Furthermore, despite the well-established role of enhancer landscapes underlying the identity of microglia, to what extent histone modifications and DNA methylation regulate microglial state switches at enhancers have not been defined. Here, using genetic fate mapping, we demonstrate the common embryonic origin of proliferative-region-associated microglia (PAM) enriched in developing white matter, and track their dynamic transitions into disease-associated microglia (DAM) and white matter-associated microglia (WAM) states in disease and aging contexts, respectively. This study links spatiotemporally discrete microglial states through their transcriptomic and epigenomic plasticity, while revealing state-specific histone modification profiles that govern state switches in health and disease.

## INTRODUCTION

Microglia are the tissue-resident macrophages in the parenchyma of the central nervous system (CNS), constantly surveying for and responding to a wide range of brain perturbations.^1–3^ In disease, microglia can present antigens, produce immunogenic cytokines, and phagocytose injured cells or disease-specific debris.^4,5^ Recently, it has been increasingly appreciated that microglia are also particularly active during development, engaging in synaptic pruning and myelinogensis.^6–9^ Modern advances in genetic labelling and single-cell RNA sequencing (scRNA-seq) technologies have allowed for the identification of distinct microglial subpopulations across these different brain contexts, including proliferative-region-associated microglia (PAM) in the developing white matter of the early postnatal brain, disease-associated microglia (DAM) in neurodegenerative disease, and white matter-associated microglia (WAM) in aging, which, together with other microglial states, contribute to the multifaceted roles of microglia in health and disease.^10–18^ While these microglial states display context-dependent gene expression patterns, they also share a core gene signature, including the upregulation of neurodegenerative disease risk genes, *Trem2* and *Apoe*, and an immune receptor gene, *Clec7a*, among others.^13,17,19^ It is unknown whether these microglial subpopulations represent ontogenically distinct lineages of microglia, or they are manifestations of plastic changes of microglial states downstream of some converging signals. Furthermore, the molecular mechanisms regulating microglial state switches remain elusive.

The enhancer repertoire underlies the transcriptional regulatory framework that governs complex gene expression programs. Enhancers are short *cis*-regulatory regions of DNA that can recruit transcription factors and interact with promoters to drive transcription.^20,21^ Enhancer activity is measured by the presence or absence of epigenetic marks including histone post-translational modifications (PTMs) and DNA methylation.^22–24^ It is well established that enhancer landscapes play a crucial role in shaping the cellular identities of tissue macrophages.^25–28^ However, how dynamic enhancer activity directs the formation of distinct microglial states in response to various microenvironmental cues is not known.

Previously, we have developed an inducible driver line, Clec7a-CreER^T2^, which allows specific labeling and tracking of *Clec7a+* microglial subsets such as PAM and DAM in different mouse models.^17^ Using cuprizone-induced demyelination, we have demonstrated the plastic changes of DAM, which are associated with white matter injury and return to homeostasis during recovery. The *in vivo* plasticity of microglial states across different contexts and over longer timescales has not been defined. Here, we systematically dissect the lineage relationships between specific states of microglia in development, aging and disease to uncover the plastic nature of microglia. In addition, we use highly sensitive epigenetic profiling approaches, CUT&Tag^29^ and whole genome bisulfite sequencing (WGBS), to characterize the dynamic changes of state-specific enhancers and predict their potential binding transcription factors that govern microglial state switches. These findings provide key insights into the molecular mechanisms underlying transcriptomic and functional heterogeneity of microglia across dynamic brain contexts.

## RESULTS

### Origin and dynamic changes of PAM during development

Previously, through single-cell RNA sequencing (scRNA-seq), we have identified proliferative-region-associated microglia (PAM) that are enriched in the corpus callosum and cerebellar white matter regions during the early postnatal development.^10^ PAM upregulate a cassette of genes, including *Clec7a*, *Itgax*, *Spp1*, *Gpnmb*, *Cd68*, compared to homeostatic microglia, and they display amoeboid morphology and are highly phagocytic. Notably, some of these marker genes and functional properties are reminiscent of peripheral myeloid cells, raising the question of whether PAM are derived from the yolk sac precursors of microglia or perinatal infiltration of peripheral immune cells.^30–32^ To determine the exact origin of PAM, we employed the Runx1-CreER^T2^ fate mapping model, and pulsed tamoxifen at embryonic day 7 (E7), which has been shown to capture the embryonic precursors leading to labeling of about 30% of total microglia across the adult brain.^31,33^ Using the same regimen, we performed immunohistochemistry on mice at postnatal day 7 (P7), when PAM were most abundant. We observed robust labeling of PAM in the developing white matter regions as well as homeostatic microglia randomly distributed in other areas of the brain (**Figure 1A, Figure S1A-C**). The labeling efficiencies for PAM and homeostatic microglia were comparable (**Figure 1B**). We saw a similar labeling pattern with another fate mapping line, Cdh5-CreER^T2,34^ which marks hemogenic endothelial cells in the early embryos (**Figure S1D**). These data suggest that PAM share the same embryonic yolk sac origin with other microglial populations, and they represent a distinct state associated with the white matter and neurogenic niches.

**Figure 1.**
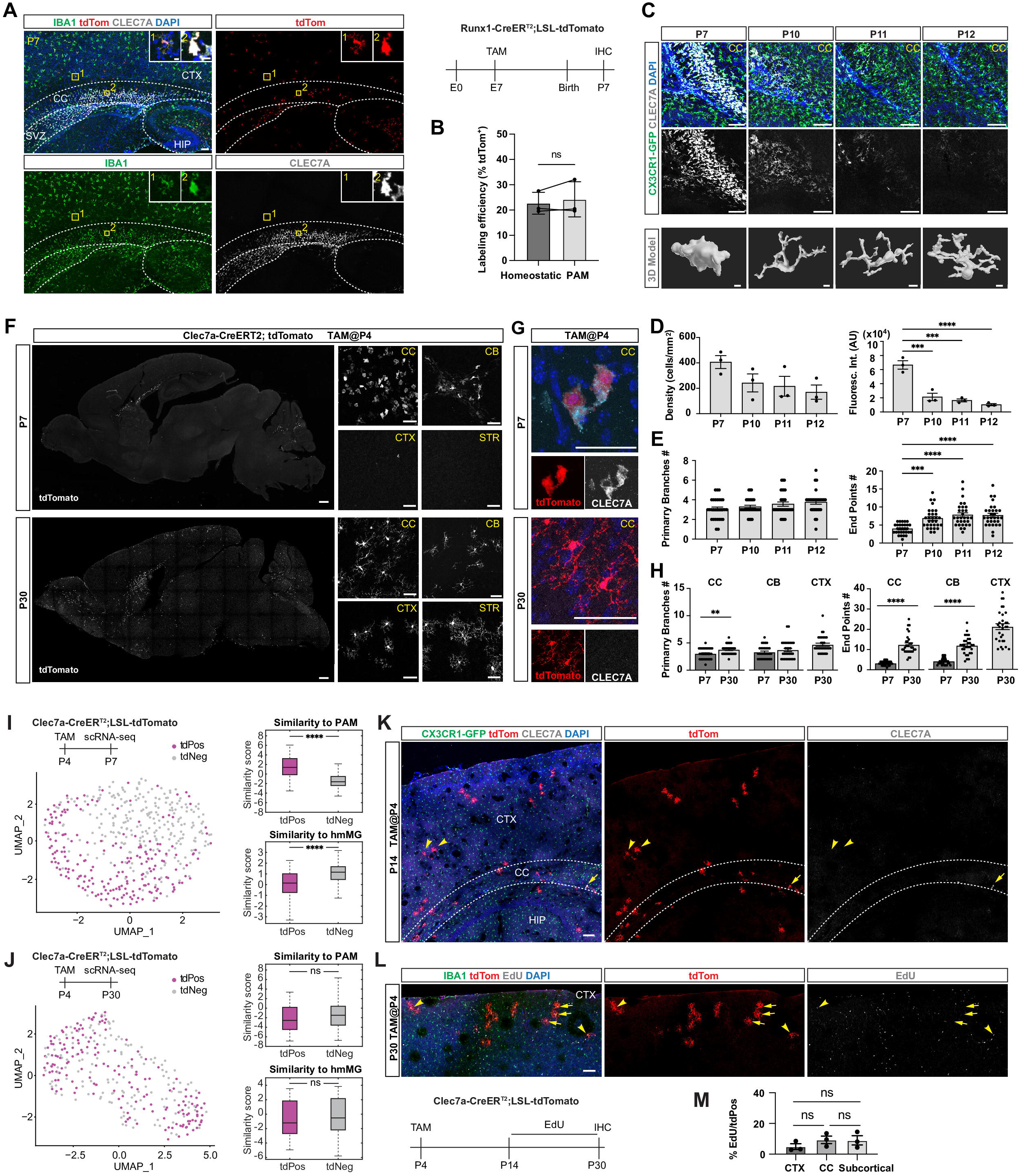
Embryonic origin and dynamic changes of PAM during development. **(A)** Representative immunostaining images showing tdTomato+ microglia (IBA1^+^) overlap with CLEC7A+ signals in the corpus callosum of a P7 mouse brain in the Runx1-CreER^T2^; LSL-tdTomato lineage tracing model. TdTomato+ microglia (IBA1^+^) in the cortical regions do not express CLEC7A. Boxed inserts showing zoomed-in images of microglia in numbered areas. Tamoxifen regimen is shown on the right. CC, corpus callosum; CTX, cortex; SVZ, subventricular zone; HIP, hippocampus. Scale bars, 100um for large images and 10um for inserts. **(B)** Quantification of labeling efficiency (percentage of tdTomato+ microglia) for microglia in cortical regions (Homeostatic) and the corpus callosum (PAM, i.e. CLEC7A+) in the Runx1-CreER^T2^; LSL-tdTomato lineage tracing model. Student’s t-test. ns, not significant. *n* = 3 mice. Error bars represent mean ± SEM. **(C)** Representative images showing changes of CLEC7A expression and morphology in PAM across developmental timepoints (P7 to P12). 3D reconstruction of microglia at each postnatal age shown at the bottom. CC, corpus callosum. Scale bars, 100um for the top panels and 5um for the bottom panels. **(D)** Quantification of CLEC7A+ microglia density (the number of cells/mm^2^) and CLEC7A fluorescence intensity (AU) for P7, P10, P11, and P12 mice. *n =* 3 mice/per group. One-way ANOVA with Tukey’s multiple comparisons test. *** *p <* 0.001, **** *p* <0.0001. Error bars represent mean ± SEM. **(E)** Quantification of microglia morphology using the numbers of primary branches (left) and endpoints (right) for P7, P10, P11, and P12 mice. *n =* 10 cells/per animal, 3 animals/per group. One-way ANOVA with Tukey’s multiple comparisons test. *** *p <* 0.001, **** *p* <0.0001. Error bars represent mean ± SEM. **(F)** Representative immunostaining images of tdTomato+ microglia in P7 (top left) and P30 (bottom left) mouse brains of the Clec7a-CreER^T2^; LSL-tdTomato mouse line after tamoxifen injection at P4. Right panels showing higher magnification images in CC, corpus callosum; CB, cerebellum; CTX, cortex; STR, striatum. Scale bars, 500um for left panels and 50um for right panels. **(G)** Representative immunostaining images showing that amoeboid tdTomato+ microglia overlap with CLEC7A signals in the CC, corpus callosum, of a P7 mouse brain after tamoxifen injection at P4. At P30, tdTomato+ microglia in the corpus callosum do not express CLEC7A and they are ramified. Scale bars, 50um. **(H)** Quantification of microglia morphology using the numbers of primary branches (left) and endpoints (right) in CC, corpus callosum and CB, cerebellum for P7 and P30 mice and CTX, cortex for P30 mice. *n =* 10 cells/per animal, 3 animals/per group. Student’s t-test. ** *p <* 0.01, **** *p* <0.0001. Error bars represent mean ± SEM. **(I)** UMAP plot showing separation of tdTomato+ and tdTomato-microglia at P7, following tamoxifen injection at P4. Box plots on the right showing the similarity scores of each group compared to PAM (top) and homeostatic microglia (hmMg; bottom). **** *p <* 0.0001. **(J)** UMAP plot showing that tdTomato+ and tdTomato-microglia are intermingled at P30, following tamoxifen injection at P4. Box plots on the right showing the similarity scores of each group compared to PAM (top) and homeostatic microglia (hmMg; bottom). ns, not significant. **(K)** Representative immunostaining images showing tdTomato+ microglia in CC, corpus callosum, and CTX, cortex of a P14 mouse brain, with tamoxifen pulsed at P4, in the Clec7a-CreER^T2^; LSL-tdTomato; CX3CR1-GFP mouse line. Yellow arrowheads point to tdTomato+ (CLEC7A-) microglia residing in intermediate cortical regions. The yellow arrow points to a rare microglial cell that still expresses CLEC7A in the corpus callosum. HIP, hippocampus. Scale bars, 100um. **(L)** Representative immunostaining images showing both tdTomato+EdU+ (arrowheads) and tdTomato+EdU-(arrows) microglia in the cortical region of a P30 brain in the Clec7a-CreER^T2^; LSL-tdTomato mouse. Tamoxifen and EdU injection regimens are shown below. Scale bars, 100um. **(M)** Quantification of tdTomato+ microglial proliferation (percentage of tdTomato+EdU+/tdTomato+) in CTX, cortex, CC, corpus callosum, and subcortical regions from P14 to P30. *n =* 3 mice. One-way ANOVA with Tukey’s multiple comparisons test. ns, not significant. Error bars represent mean ± SEM. **See also Figure S1.**

To characterize the dynamic changes of PAM in the first two postnatal weeks, we performed time course analyses on their density, morphology and marker gene expression. Interestingly, within only three days following the peak appearance (from P7 to P10), PAM sharply downregulated the marker gene *Clec7a*, and they also became more ramified, while their density remained largely unchanged (**Figure 1C-E**). By P14, PAM were almost undetectable based on marker gene expression in the corpus callosum.^10^ This disappearance of PAM may be due to cell death (e.g. apoptosis, ferroptosis) or a state change with the downregulation of their signature genes.^17,30,35^ To distinguish these two possibilities, we leveraged our recently published driver line, Clec7a-CreER^T2^, which specifically labels PAM (and other *Clec7a+* microglial states in disease settings) in the developing white matter and thus allow us to follow the fate of PAM into later timepoints.^17^ By pulsing tamoxifen at P4, we observed CLEC7A+ microglia co-labeled with the tdTomato reporter in the corpus callosum and cerebellar white matter at P7 as expected (**Figure 1F and 1G**). When examining the tissues at P30, we found that numerous tdTomato+ microglia were still present in the PAM-occupying regions, suggesting that the PAM lineage cells survived into adulthood. Interestingly, these tdTomato+ microglia also appeared in other parts of the brain, such as upper cortical layers and striatal regions (**Figure 1F**). In addition, the tdTomato+ microglia in P30 exhibited ramified morphology and were stained negative for CLEC7A (**Figure 1G, 1H**). These data suggest that PAM lose marker gene expression and adopt homeostatic features following the early postnatal development.

To examine the changes of PAM at the transcriptomic level, we performed scRNA-seq on tdTomato+ and tdTomato-microglia isolated from P7 and P30 brains (**Figure 1I, 1J**). As expected, P7 tdTomato+ microglia, representing PAM, were separated from tdTomato-cells, which mainly contained homeostatic microglia, on the UMAP (**Figure 1I**). In contrast, these two groups of cells were no longer separable at P30, suggesting global conversion of PAM into homeostatic microglia in the adult brain (**Figure 1J**). We will refer to these PAM-derived homeostatic microglia as exPAM hereafter.

We were curious about the regional distribution of exPAM, particularly those found in the upper cortical layers (**Figure 1F**) and wanted to understand how they arrived there. Although we could not monitor PAM in real time due to their deep anatomical location and the long duration of potential changes, we reproducibly found incidences of exPAM between the corpus callosum and superficial layers of the brain at intermediate timepoints (e.g. P14), indicating cell migration (**Figure 1K**). To assess the involvement of microglial proliferation, we tracked cell divisions with EdU injections on top of PAM lineage tracing (**Figure 1L**). Despite wide-spread EdU labeling across the entire brain, especially along the rostral migratory stream, only less than 10% tdTomato+ cells were co-labeled with EdU (**Figure 1M, Figure S1E**). These data suggest that a combination of migration and limited proliferation contribute to exPAM outside the white matter regions. Taken together, this series of fate mapping experiments illustrate PAM as a dynamic microglial state that arises from yolk sac precursors and converts to homeostatic microglia across multiple brain regions later in life.

### Plasticity of PAM during injury and aging

To obtain a more comprehensive view about the plasticity of PAM, we wanted to first analyze the regional distribution of exPAM in a quantitative manner. We generated Clec7a-CreER^T2^; LSL-tdTomato; CX3CR1-GFP dual-color labeling mice and used the fluorescence reporters to visualize PAM or exPAM (in tdTomato) and all microglia (in GFP) following whole brain tissue clearing.^36^ Consistent with immunostaining on tissue sections, PAM were mainly restricted to the corpus callosum, certain white matter tracks (e.g. cingulum bundle, optic radiation), and deep cortical regions (e.g. primary motor and somatosensory areas in layer 6) (**Figure 2A-C, Figure S2A-B, Videos S1-S2, Table S1**). While exPAM in the P30 brain were still present in the white matter regions, their density was markedly reduced (**Figure 2C, Figure S2B**). In addition, exPAM were also specifically enriched in primary somatosensory layer 1 and certain layer 2/3 cortical regions, as well as in amygdalar and thalamus-related nuclei (**Figure 2B-C, Figure S2B, Videos S3-S4, Table S1**). The reproducibility of this labeling pattern suggests controlled microglial migratory routes possibly due to functional recruitment of exPAM to these brain regions and/or anatomical scaffolds facilitating cell movement.

**Figure 2.**
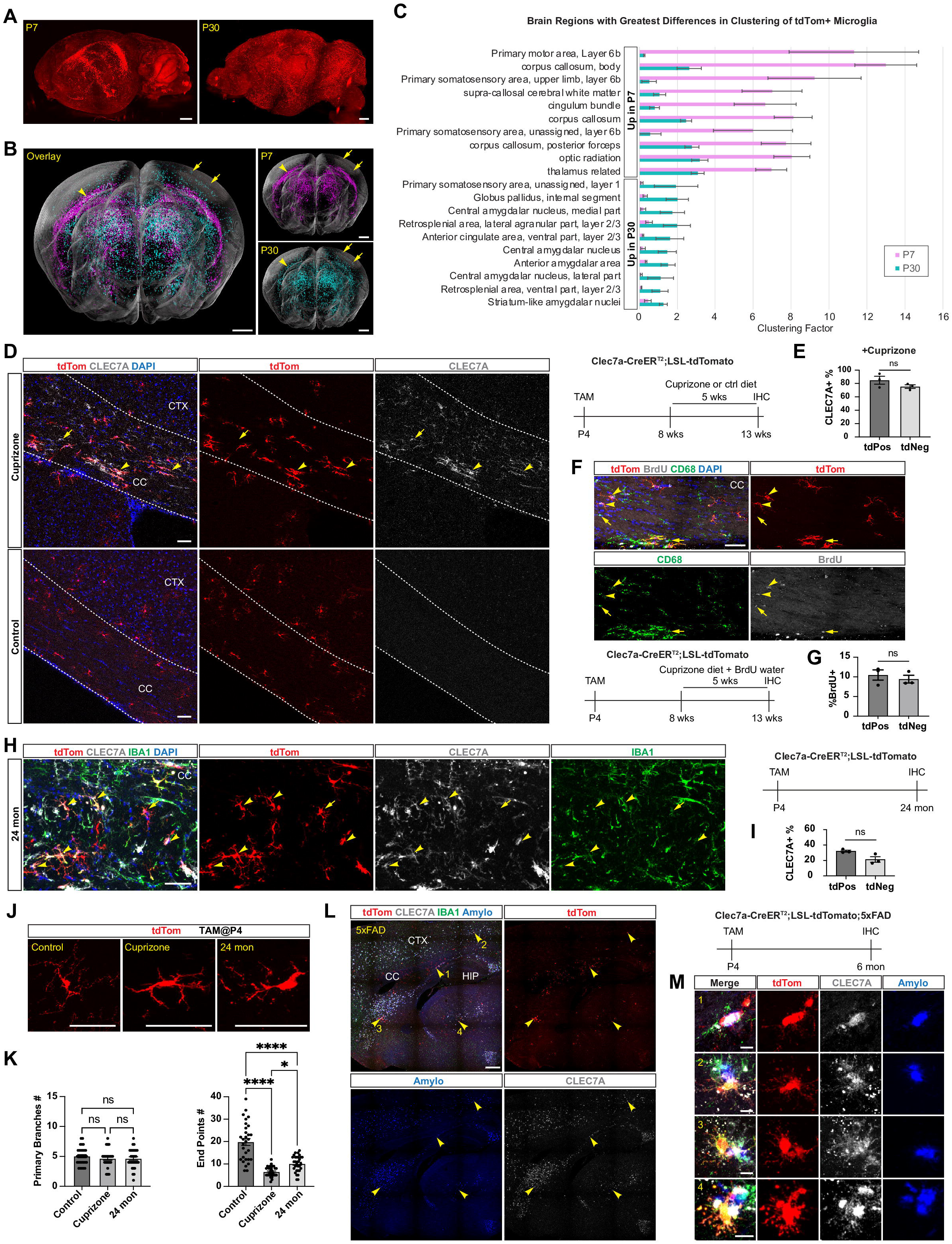
Plasticity of PAM during injury and aging. **(A)** Lateral view of a cleared whole brain of Clec7a-CreER^T2^; LSL-tdTomato; CX3CR1-GFP mice at P7 (left) and P30 (right) after tamoxifen injection at P4. Only the tdTomato signals are shown. Scale bars, 1000um. **(B)** Coronal view of tdTomato+ microglia from cleared whole brains projected onto a standard brain model. P7 microglia are shown in magenta and P30 microglia are shown in cyan, with the overlay image shown on the left. The arrowhead points to the corpus callosum region where both P7 and P30 tdTomato+ microglia are found. Arrows highlight tdTomato+ microglia in the superficial cortical layers of a P30 brain. Scale bars, 1000um. **(C)** Quantification of brain regions with greatest differences in cell clustering of tdTomato+ microglia from P7 (magenta) and P30 (cyan) brains. The regions that have higher enrichment for P7 microglia are shown on the top and the regions with higher enrichment for P30 microglia are shown on the bottom. The bars are ordered by fold differences for each time point. Error bars represent mean ± SE. **(D)** Representative images showing tdTomato+ microglia co-stained with CLEC7A antibodies in the CC, corpus callosum, of cuprizone-treated (top) and control (bottom) Clec7a-CreER^T2^; LSL-tdTomato mice after tamoxifen injection at P4. Tamoxifen injection and cuprizone diet regimens are shown on the right. TdTomato+ microglia that express CLEC7A are indicated by arrowheads. The yellow arrow points to a microglial cell that expresses CLEC7A+ but is not labeled by tdTomato. TdTomato+ from the control mice do not express CLEC7A. CTX, cortex. Scale bars, 50um. **(E)** Quantification of the percentage of CLEC7A+ microglia in tdTomato+ (percent tdTomato+CLEC7A+/tdTomato+) and tdTomato-(percent tdTomato-CLEC7A+/ tdTomato-) microglia (all IBA1+) in cuprizone-treated mice, following tamoxifen injection at P4. *n* = 3 mice. Student’s t-test. ns, not significant. Error bars represent mean ± SEM. **(F)** Representative images showing overlaps between tdTomato+CD68+ microglia (PAM-derived DAM) and BrdU signals in CC, corpus callosum, of Clec7a-CreER^T2^; LSL-tdTomato mice after tamoxifen injection at P4 and 0.2% cuprizone treatment for 5 weeks (P60 onwards). Arrowheads indicate tdTomato+CD68+BrdU+ microglia, and arrows point to tdTomato-CD68+BrdU+ microglia (non-PAM-derived DAM). Tamoxifen and cuprizone regimens are shown below. Scale bars, 50um. **(G)** Quantification of percentage of BrdU+ microglia in tdTomato+ (PAM-derived) and tdTomato-(non-PAM-derived) microglia that are also CD68+ (DAM). *n* = 3 mice. ns, not significant. Error bars represent mean ± SEM. **(H)** Representative images showing tdTomato+CLEC7A+ microglia (arrowheads) in CC, corpus callosum of 24-month-old Clec7a-CreER^T2^; LSL-tdTomato mice after tamoxifen injection at P4 (regimen shown on the right). The yellow arrow points to a tdTomato^+^ microglial cell that does not express CLEC7A. Scale bars, 50um. **(I)** Quantification of the percentage of CLEC7A+ microglia in tdTomato+ (percent tdTomato+CLEC7A+/tdTomato+) and tdTomato-(percent tdTomato-CLEC7A+/ tdTomato-) microglia (all IBA1+) in 24-month-old Clec7a-CreER^T2^; LSL-tdTomato mice after tamoxifen injection at P4 (regimen shown on the right). *n* = 3 mice. Student’s t-test. ns, not significant. Error bars represent mean ± SEM. **(J)** Representative higher-magnification images of tdTomato+ microglia in control (left), cuprizone-treated (middle), and 24-month-old mice (right) showing morphological differences. Tamoxifen injection is done at P4 for all conditions. Scale bars, 50um. **(K)** Quantification of tdTomato+ microglia morphology using the numbers of primary branches (left) and endpoints (right) in control, cuprizone-treated, and 24-month-old mice. Tamoxifen injection is done at P4 for all conditions. *n* = 10 cells/per animal, 3 animals/per group. One-way ANOVA with Tukey’s multiple comparisons test. ns, not significant. * *p <* 0.05, **** *p <* 0.0001. Error bars represent mean ± SEM. **(L)** Representative images showing tdTomato+ microglia co-stained with CLEC7A and amyloid plaques (Amylo-Glo) in 6-month-old Clec7a-CreER^T2^; LSL-tdTomato; 5xFAD mice after tamoxifen injection at P4 (regimen shown on the right). Numbered arrowheads indicate tdTomato+ microglia that express CLEC7A around amyloid plaques. CC, corpus callosum; CTX, cortex; HIP, hippocampus. Scale bars, 500um. **(M)** Higher-magnification images of the numbered areas in (L) showing tdTomato+CLEC7A+ microglia around amyloid plaques in the 5xFAD model. Scale bars, 10um. **See also Figure S2, Table S1, Videos S1-S4.**

Next, we wanted to understand whether exPAM can become reactivated upon a stimulus later in life. We decided to focus on their response in white matter injuries. We injected tamoxifen in the Clec7a-CreER^T2^; LSL-tdTomato reporter mice at P4 and fed the adult mice with either control or cuprizone diet to generate demyelination. Indeed, about 80% of tdTomato+ microglia were stained positive for CLEC7A in the corpus callosum of the cuprizone-treated mice, whereas similarly labeled cells in the control condition remained negative for this reactive marker (**Figure 2D and 2E**). The CLEC7A+ microglia also expressed CD11c and CD68, suggesting that they assumed the disease-associated microglia (DAM) reactive phenotype (**Figure S2C**). Consistently, these cells displayed less ramified morphology (**Figure 2J and 2K**). Interestingly, a subset of tdTomato-microglia in the cuprizone-treated group also became reactive, and there seemed to be no preference between the tdTomato+ (exPAM) and tdTomato-cells in terms of the generation of DAM upon the white matter injury (**Figure 2E**). To discern the extent by which exPAM, as opposed to their daughter cells from cell proliferation, directly contributed to DAM, we provided BrdU drinking water to the reporter mice throughout the cuprizone treatment (**Figure 2F**). We found that the vast majority (about 90%) of tdTomato+CLEC7A+ microglia in the white matter were never labeled by BrdU, suggesting minimal cell proliferation (**Figure 2G**). These data underscore the plasticity of reactive microglia as they undergo state switches from PAM to exPAM (homeostasis) and to DAM in a white matter disease later in life (**Figure S2E**). Can exPAM become reactivated during a natural process such as normal aging? Previously, white matter-associated microglia (WAM), which share a similar gene signature with PAM (and DAM), have been identified in the aging brain,^14^ prompting us to assess their lineage relationship. Surprisingly, by rearing the P4 injected reporter mice for additional two years, we found that numerous tdTomato+ labeled microglia remained in the white matter regions (**Figure 2H**). About 30% of these cells turned on reactive markers and simultaneously exhibited reactive morphology, suggesting a state switch to WAM (**Figure 2H-K, Figure S2D**). Although it is not feasible to track cell proliferation over such a long period of time, given the slow turnover of microglia,^37–39^ we speculate that some exPAM may directly contribute to the pool of WAM in aging.

Up until this point, our analyses have been focusing on the white matter where these reactive microglia populations reside. We wondered to what extent exPAM can generate DAM in a disease with primary pathology outside the white matter. To address this question, we crossed the Clec7a-CreER^T2^; LSL-tdTomato reporter mice to 5xFAD, an amyloidosis model of Alzheimer’s disease (AD).^40^ In 6-month-old 5xFAD mice, amyloid plaques were primarily built up in the cortex, hippocampus and striatum, with little accumulation in the white matter (**Figure 2L**). Interestingly, despite the presence of plaques in the areas adjacent to the corpus callosum, the regional distribution of exPAM largely remained the same as seen in the healthy adult brain (**Figure 1F, 2L**). As such, only 3.0±0.7% of exPAM that directly surrounded plaques became reactive in response to AD pathology (**Figure 2L and 2M**), which is consistent with the notion that microglial cell bodies are mostly stationary in the adult brain.^2,3^ Nonetheless, this observation highlights that exPAM are fully capable of responding to amyloid deposition and switching to the DAM state when they physically interact with plaques in AD.

Together, these results demonstrate the extraordinary plasticity of exPAM, which undergo state switches to DAM or WAM in response to disease or during normal aging.

### Epigenomic profiling of PAM and identification of state-specific gene regulatory elements

Given the plasticity of PAM, we wanted to understand the molecular mechanisms that regulate microglial state switches. It has been shown that enhancer landscapes play a critical role in regulating the identities of tissue macrophages including microglia.^25–27^ However, epigenomic features, such as histone modifications and DNA methylomes, underlying specific microglial states have not been defined. Due to the rarity of these reactive microglial subpopulations in their respective brain contexts, we reasoned that conventional chromatin profiling methods would be insufficient to capture the dynamic signals of chromatin factors to readily identify *de novo* gene regulatory elements active in PAM and DAM. We therefore performed Cleavage Under Targets and Tagmentation (CUT&Tag) for H3 lysine 27 acetylation (H3K27ac) and H3 lysine 4 monomethylation (H3K4me1), canonical histone marks associated with active gene regulatory elements (**Figure 3A**).^29,41^ As a proof of concept, we assessed differential H3K27ac and H3K4me1 signals at known microglial versus myeloid promoters and enhancers using as few as 2,000 wildtype microglia. Indeed, aggregate analysis revealed robust activation of the promoters and enhancers associated with microglia-specific genes but not myeloid-specific genes^25,42^ as measured by these histone marks (**Figure 3B**). When we examined H3K27ac and H3K4me1 signals as a function of gene expression, we observed strong concordance between the levels of these marks and gene expression at promoters and enhancers. Therefore, CUT&Tag on small numbers of microglia can specifically profile H3K27ac and H3K4me1 signals as an accurate proxy of promoter and enhancer activity that is tightly linked to gene expression.

**Figure 3.**
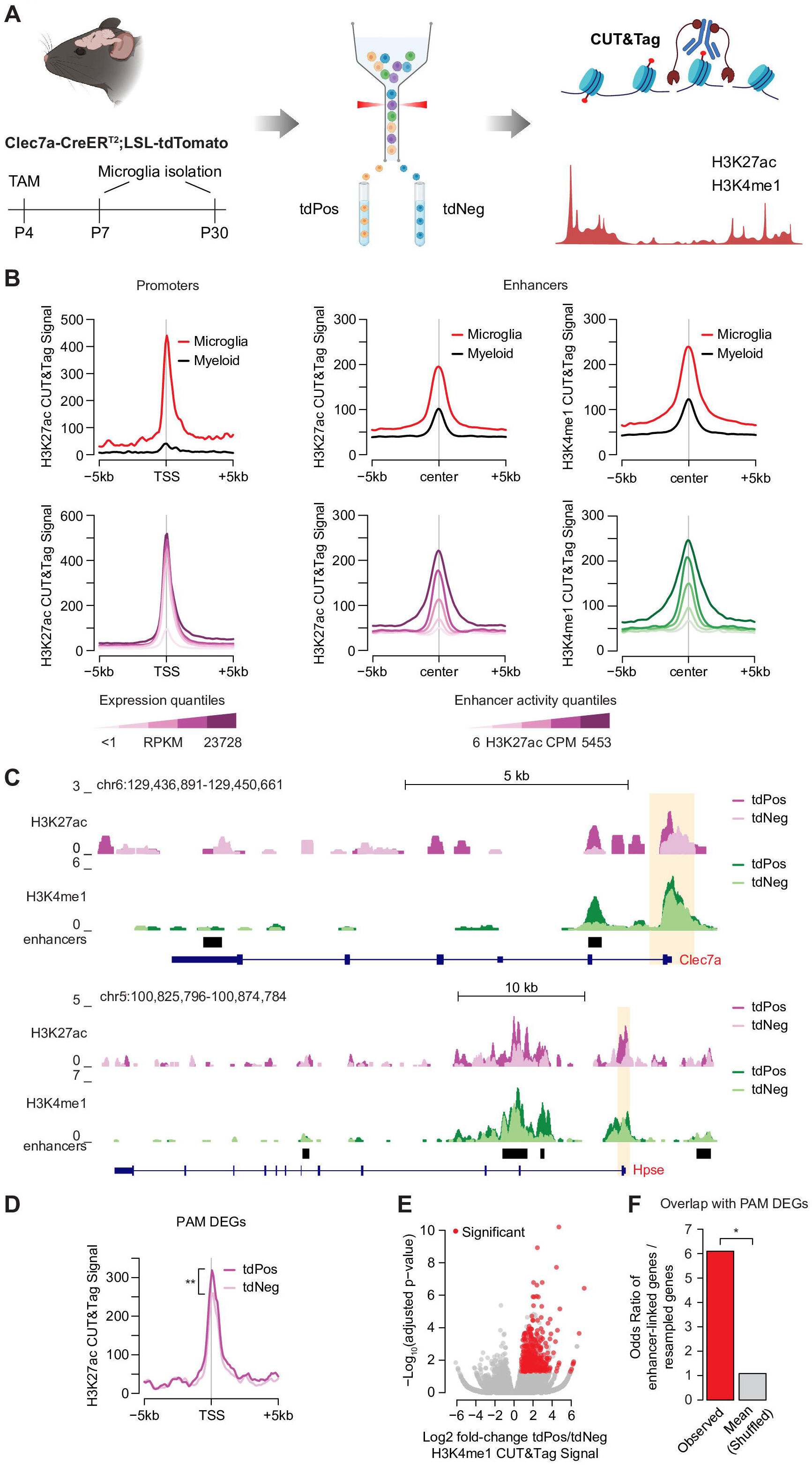
Identification of putative enhancers that regulate PAM state. **(A)** Schematic depiction of experimental design for CUT&Tag experiments in P7 and P30 brains. TAM, tamoxifen. **(B)** Aggregate H3K27ac signals at microglial gene promoters versus myeloid gene promoters (top left) and across quantiles of microglial gene expression (bottom left). Aggregate H3K27ac and H3K4me1 signals at microglial enhancers versus myeloid enhancers, and across quantiles of microglial enhancer H3K27ac levels (right). Microglia and myeloid gene expression and enhancer data were obtained from published work.^25,42^ **(C)** Genome browser view of CUT&Tag data for tdPos and tdNeg microglia isolated from P7 at two example gene loci, *Clec7a* and *Hpse*, both upregulated in PAM. Promoter regions are highlighted in orange. Enhancer regions (marked by black rectangular shapes) were defined by identifying H3K4me1 peaks using the MACS2 peak calling algorithm and filtering out peaks overlapping with gene promoter regions (1kb around annotated TSS).^71^ **(D)** Aggregate H3K27ac signals at gene promoter regions of PAM upregulated DEGs (Log2 fold-change >1.0, FDR <0.05). PAM gene expression data were obtained from published work.^10^ ** *p* < 0.01; paired t-test across mean normalized CPM values between tdPos and tdNeg microglia bioreplicates (n = 4). **(E)** Volcano plot of fold-changes in H3K4me1 levels in tdPos versus tdNeg microglia at P7. Enhancers were called as significantly more active in the tdPos versus tdNeg if they had significant upregulation of H3K4me1 signal (FDR <0.05) and had concordant enrichment in H3K27ac signal (Log2 fold-change >0). **(F)** Odds ratio of overlap of genes associated with PAM enhancers by GREAT analysis and PAM DEGs compared to odds ratio of overlap of expression-matched resampled genes (mean of 1000 resamples). * *p* <0.05; Fischer’s exact test.

Having validated our epigenomic profiling methods, we sought to identify gene regulatory elements that control the PAM state. We collected CD45^int^CD11b^+^tdTomato^+^ (tdPos) and CD45^int^CD11b^+^tdTomato^-^ (tdNeg) cells, representing PAM and homeostatic microglia, respectively, from P7 brains of the Clec7a-CreER^T2^ reporter mice. Visual inspection and aggregate analysis of promoters associated with previously identified differentially upregulated genes^10^ in PAM showed higher H3K27ac signals in tdPos over tdNeg microglia (**Figure 3C, D**). To identify PAM enhancers, we first classified enhancers based on the presence of H3K4me1, and further performed edgeR differential count analysis^43^ on H3K4me1 and H3K27ac signals in tdPos microglia over tdNeg microglia. We identified 385 *de novo* enhancers with significantly enriched H3K4me1 and H3K27ac signals in the tdPos population (**Figure 3E and 4B, Table S2**). Interestingly, the increases of these histone modifications at the enhancers were more pronounced compared to those at the promoters (**Figure 3D and 4B**), consistent with the notion that enhancer activity tends to be more dynamic with gene expression changes.^28^ Analysis of gene-enhancer linkage via GREAT^44^ revealed that these enhancers were significantly associated with PAM differentially expressed genes (DEGs) (**Figure 3F**). Therefore, we have uncovered a set of enhancers that are likely to be functionally relevant in PAM.

**Figure 4.**
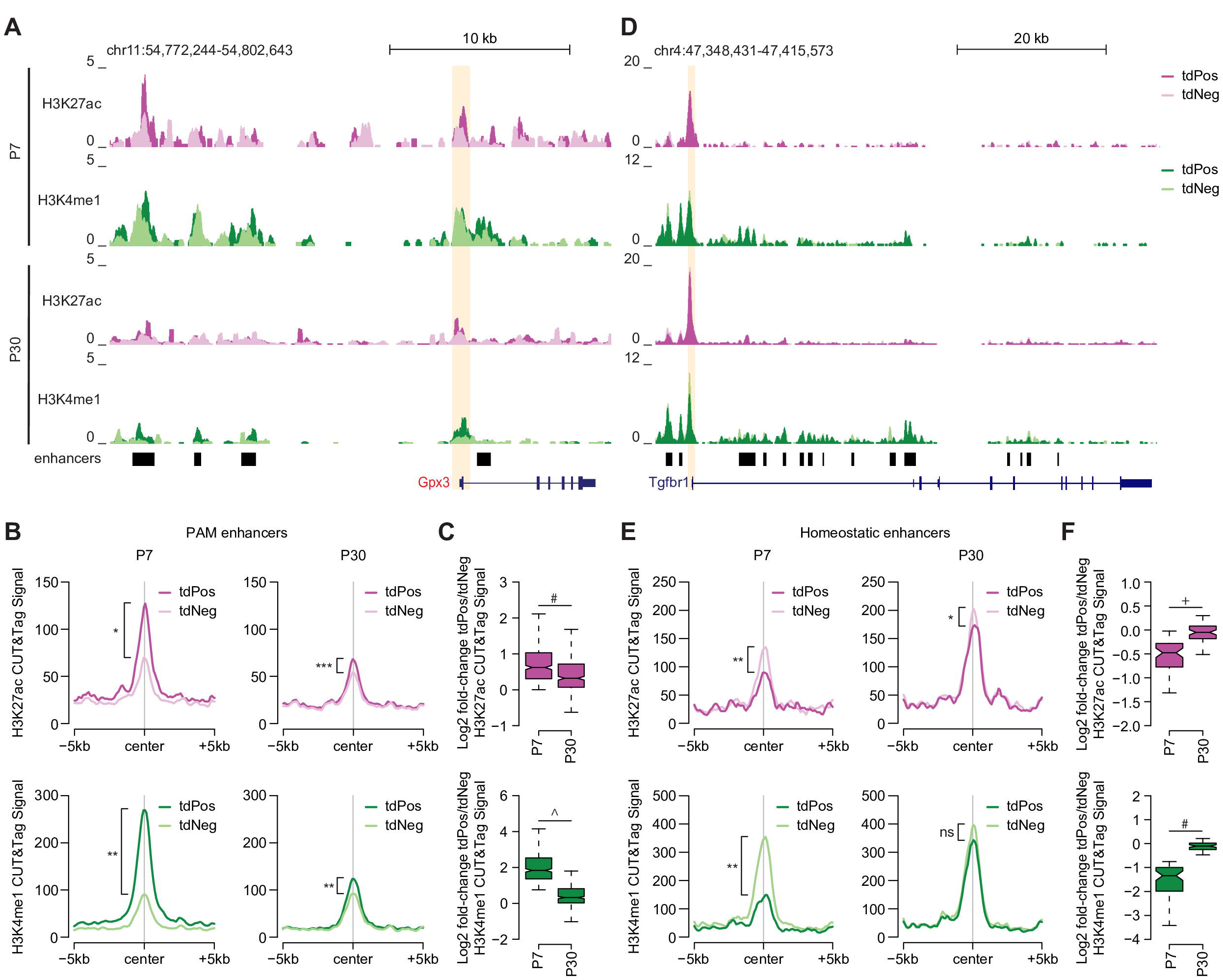
State-specific enhancers exhibit dynamic activity that reflect PAM transition to a homeostatic state in adult mouse brain. **(A)** Genome browser view of CUT&Tag data for tdPos and tdNeg microglia isolated from P7 and P30 at a PAM-specific DEG, *Gpx3*. Promoter is highlighted in orange. Enhancer regions are marked by black rectangular shapes. Overlap of nearby enhancers with enriched H3K27ac and H3K4me1 signals in the tdPos over the tdNeg population at P7 that subsequently decrease in signal at P30 illustrates PAM- associated enhancer dynamics. **(B)** Aggregate H3K27ac and H3K4me1 signals in tdPos versus tdNeg microglia at PAM enhancers at P7 and P30 timepoints. * *p* < 0.05; ** *p* < 0.01; *** *p* < 0.001; paired t- test across mean normalized CPM values between tdPos and tdNeg microglia bioreplicates (P7, n = 4; P30, n =7). **(C)** Fold-changes of H3K27ac and H3K4me1 signals in tdPos versus tdNeg microglia at PAM enhancers at P7 compared to P30. # *p* < 1e-10; ^ *p* < 1e-15; Wilcoxon test. **(D)** Genome browser view of CUT&Tag data for tdPos and tdNeg microglia isolated from P7 and P30 at a homeostatic microglial gene, *Tgfbr1*. Promoter is highlighted in orange. Enhancer regions are marked by black rectangular shapes. Overlap of nearby enhancers with similar H3K27ac and H3K4me1 signals in the tdPos and the tdNeg population at P7 that subsequently increase in signal at P30 illustrates homeostatic microglia-associated enhancers are upregulated during PAM transition to homeostasis in adulthood. **(E)** Aggregate H3K27ac and H3K4me1 signals in tdPos versus tdNeg microglia at homeostatic enhancers at P7 and P30 timepoints. * *p* < 0.05; ** *p* < 0.01; ns, not significant; paired t-test across mean normalized CPM values between tdPos and tdNeg microglia bioreplicates (P7, n = 4; P30, n =7). **(F)** Fold-changes of H3K27ac and H3K4me1 signals in tdPos versus tdNeg microglia at homeostatic enhancers at P7 compared to P30. + *p* < 1e-5; # *p* < 1e-10; Wilcoxon test. **See also Table S2.**

### Dynamic changes of histone modifications at enhancers reflect microglial state transitions

Given the transcriptomic plasticity of PAM during development (**Figure 1I and 1J**), we next wanted to investigate whether PAM-specific enhancers are involved in regulating the PAM-to-exPAM state transition. To this end, we isolated CD45^int^CD11b^+^tdTomato^+^ (tdPos) and CD45^int^CD11b^+^tdTomato^-^ (tdNeg) microglia, representing exPAM (PAM-derived homeostatic microglia) and non-PAM-derived homeostatic microglia, from P30 brains of the Clec7a-CreER^T2^ reporter mice. We performed CUT&Tag experiments and compared the results with the P7 data. Visual inspection showed that the enhancers around PAM upregulated genes (e.g. *Gpx3*) were more active in the tdPos population and they were decreased in activity by adulthood (P30) (**Figure 4A**). To obtain a comprehensive view of enhancer dynamics during this microglial state transition, we quantified the aggregated H3K27ac and H3K4me1 CUT&Tag signals at all *de novo* called PAM enhancers between tdPos and tdNeg microglia from P7 and P30 brains. Consistent with the observation for selected genes, we found a global reduction of PAM enhancer activities, particularly in the tdPos population, down to near tdNeg levels by P30 (**Figure 4B and 4C**). This data suggests that histone modifications at state-specific enhancers are highly dynamic, reflecting the transition from the PAM state to a homeostatic state.

In parallel with the identification of PAM enhancers, we were also curious about the gene regulatory elements that promote microglial homeostasis. Therefore, we performed edgeR analysis to identify enhancers with differentially enriched H3K4me1 and H3K27ac signals in tdNeg microglia compared to tdPos microglia at P7 (**Table S2**). Visual inspection of microglial homeostasis-associated genes (e.g. *Tgfbr1*) confirmed stronger signals of homeostatic enhancers in the tdNeg population (**Figure 4D**). We also quantified the aggregated changes of all *de novo* called homeostatic enhancers from P7 to P30. While these enhancers were less active in the tdPos population at P7 as expected, they robustly increased in activity by P30 becoming barely distinguishable in H3K27ac and H3K4me1 levels from the tdNeg population (**Figure 4E and 4F**). Taken together, these results point to epigenetic plasticity of microglial states, highlighting that the dynamical changes of histone modifications at state-specific enhancers underlie the microglial state switch from developmental reactivity to homeostasis.

### Identification of DAM-specific and shared enhancers with PAM state

Having established the enhancer landscapes of microglial states in physiological settings, we wanted to extend our analysis to a disease context. We decided to focus on DAM in AD, which is highly relevant to the disease progression and may have a broader impact on other neurodegenerative conditions.^45^ Despite the limited contribution of PAM to this microglial subset (**Figure 2L**), PAM and DAM (AD) share notable transcriptional, morphologic, and metabolic similarities in addition to the upregulation of certain distinct genes in each state.^10,17^ These observations have led us to ask how enhancer activity shapes the DAM state in relation to PAM.

To address this question, we administered tamoxifen to the Clec7a-CreER^T2^ reporter mice in the background of 5xFAD, which has been shown to robustly label DAM at the peak of amyloidosis.^17^ This allowed us to isolate tdPos and tdNeg microglia for epigenetic profiling of H3K27ac and H3K4me1 by CUT&Tag as we did before. Visual inspection and aggregate analysis at promoters of genes known to be upregulated in DAM^12,17^ indicated enriched H3K27ac signals in tdPos microglia compared to tdNeg microglia (**Figure 5A and 5B**). Importantly, we identified 840 significantly active enhancers in the tdPos population, which collectively showed a bigger impact on separating DAM (tdPos) versus homeostatic (tdNeg) states compared to the chromatin status at promoters (**Figure 5B and 5C, Table S2**). Again, GREAT analysis revealed that these enhancers were significantly associated with DAM DEGs (**Figure 5D**). These results suggest that we have identified a cohort of relevant enhancers for the activation of DAM in AD.

**Figure 5.**
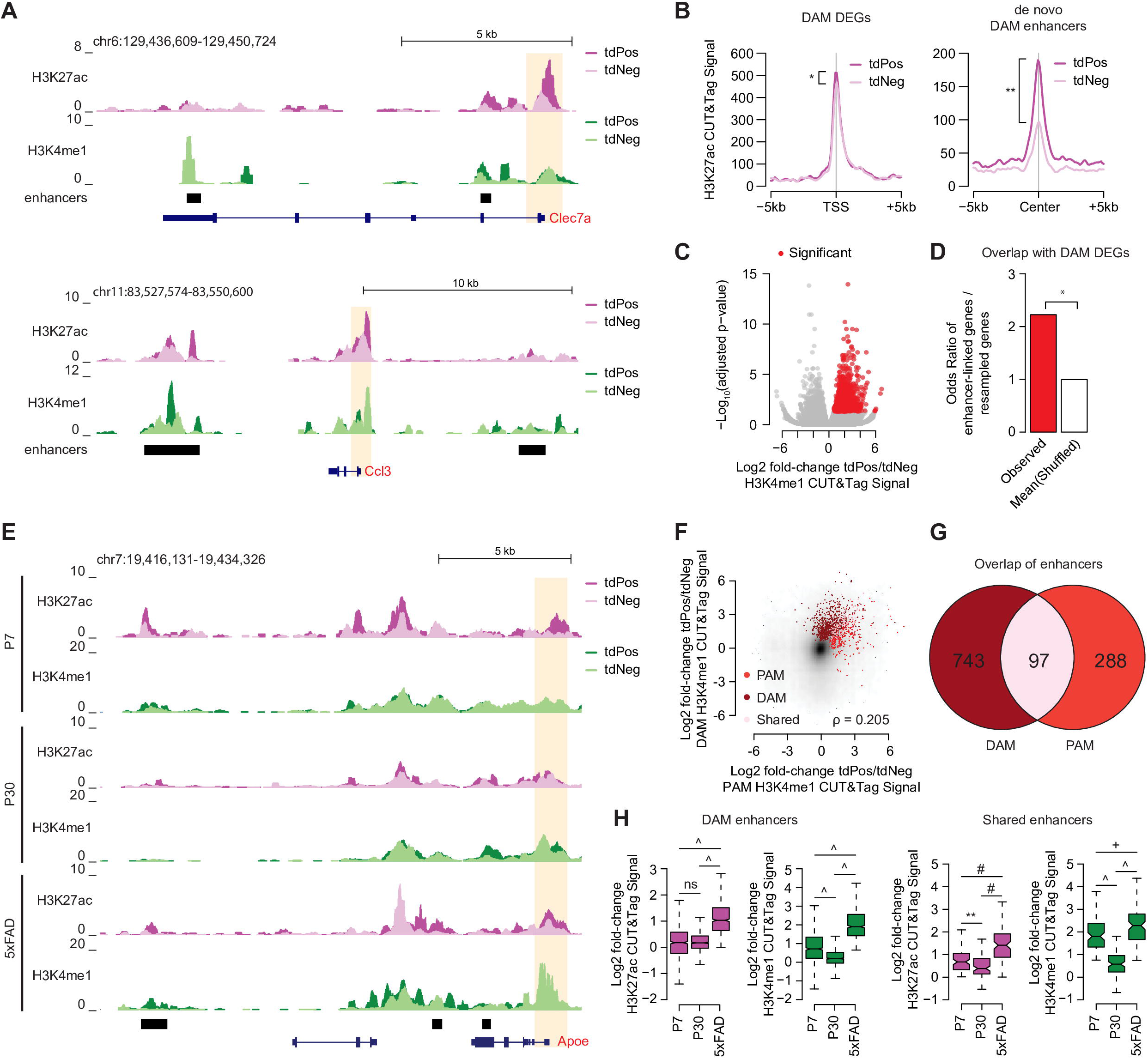
DAM show state-specific and shared enhancer activities with PAM. **(A)** Genome browser view of CUT&Tag data for tdPos and tdNeg microglia isolated from the Clec7a-CreER^T2^;LSL-tdTomato;5xFAD mouse model at two DAM DEG loci, *Clec7a* and *Ccl3*. Promoter is highlighted in orange. Enhancer regions are marked by black rectangular shapes. **(B)** Aggregate H3K27ac signals at gene promoter regions of DAM upregulated DEGs (Log2 fold-change >1.0, FDR <0.05) (left). DAM gene expression data were obtained from published work.^17^ Aggregate H3K27ac signals in tdPos versus tdNeg microglia at *de novo* called DAM enhancers in the 5xFAD model (right). * *p* < 0.05; ** *p* < 0.01; paired t-test across mean normalized CPM values between tdPos and tdNeg microglia bioreplicates (n = 5). **(C)** Volcano plot of fold-changes in H3K4me1 levels in tdPos versus tdNeg microglia in the 5xFAD model. Enhancers were called as significantly more active in the tdPos versus tdNeg if they had significant upregulation of H3K4me1 signal (FDR <0.05) and had concordant enrichment in H3K27ac signal (Log2 fold-change >0). **(D)** Odds ratio of overlap of genes associated with DAM enhancers by GREAT analysis and DAM DEGs compared to odds ratio of overlap of expression-matched resampled genes (mean of 1000 resamples). * *p* <0.05; Fischer’s exact test. **(E)** Genome browser view of CUT&Tag data for tdPos and tdNeg microglia isolated from P7 and P30 timepoints, as well as the Clec7a-CreERT2; LSL-tdTomato; 5xFAD mouse model at a shared PAM and DAM upregulated DEG, *Apoe*. Promoters are highlighted in orange. Enhancer regions are marked by black rectangular shapes. The leftmost enhancer shows elevated signals for H3K27ac and H3K4me1 in tdPos microglia from P7 and 5xFAD conditions, but the signals are diminished at P30. **(F)** Comparison of changes in H3K4me1 at enhancers in tdPos versus tdNeg microglia from P7 with H3K4me1 changes at the same enhancers in tdPos versus tdNeg microglia isolated from the 5xFAD model. PAM-specific, DAM-specific, and shared PAM and DAM enhancers (edgeR, FDR <0.05) are annotated. **(G)** Venn diagram showing the numbers of specific and overlapping active enhancers in PAM and DAM (tdPos) relative to condition-specific homeostatic cells (tdNeg). **(H)** Fold-changes of H3K27ac and H3K4me1 signals in tdPos versus tdNeg microglia at DAM-specific and PAM/DAM shared enhancers across a series of conditions (P7, P30, and 5xFAD model). ** *p* < 0.01; + *p* < 1e-5; # *p* < 1e-10; ^ *p* < 1e-15; ns, not significant; Wilcoxon test.

Given that DAM and PAM share a core gene signature, we wanted to investigate whether they also share an underlying epigenetic profile. Using *Apoe* as an example, which is the biggest genetic risk gene for late-onset AD and one of the most upregulated genes in PAM and DAM, we found specific enrichment of H3K27ac and H3K4me1 signals at a downstream distal enhancer in the tdPos populations of P7 and 5xFAD brains, suggesting a possible shared role of this regulatory element in these two conditions (**Figure 5E**). Cross correlation analysis revealed that enhancers significantly active in PAM tended to also be active in DAM, and vice versa (**Figure 5F**). Consistently, there was a substantial overlap between *de novo* called PAM and DAM enhancers (**Figure 5F and 5G**). Quantitative analysis demonstrated that the shared enhancers were not only active at P7 and in the 5xFAD model but were also deactivated at P30 during a predominantly microglial homeostatic context (**Figure 5H**). Together, these results suggest that PAM and DAM have distinct and shared epigenetic properties that may impact their respective contributions to development and disease.

### Stable DNA hypomethylation is permissive for microglial activation

Histone modifications can emerge and disappear rapidly, within minutes to hours, making them well-suited for regulating microglial activation over short timescales.^46^ On the other hand, DNA methylation at CpG dinucleotides (mCG) is another type of epigenetic regulation, which involves covalent modifications on DNA and has much slower turnover rates (ranging from hours, days to years depending on the contexts).^47,48^ We were curious whether such a relatively stable form of epigenetic mechanisms plays a role in regulating microglial state switches. To address this question, we first performed whole genome bisulfite sequencing (WGBS) to measure mCG levels at base pair resolution from adult wildtype microglia. Aggregate analysis of mCG levels at microglia- and myeloid-specific gene regulatory elements revealed greater depletion of mCG at microglial promoters and enhancers, as expected (**Figure S3**). Furthermore, we observed a robust correlation between mCG depletion and gene expression at enhancers compared to promoters which exhibited a more binary relationship between mCG loss and gene expression (**Figure S3**). These results highlight that the degree of dynamic mCG patterning at enhancers is associated with finetuning gene expression in microglia.

To examine the extent by which DNA methylomes underlie microglial reactivity, we isolated tdPos and tdNeg microglia from P7, P30 and 5xFAD mice, followed by WGBS (**Figure 6A**). Interestingly, regardless of the stages and conditions, both tdPos (PAM in P7, exPAM in P30, or DAM in 5xFAD) and tdNeg (homeostatic microglia) populations displayed similar levels of mCG depletion at the promoter and enhancer regions of reactive marker genes (e.g. *Clec7a*, *Trem2*), as well as homeostatic genes (e.g. *Tgfbr1*) (**Figure 6B and 6E**). Focusing on the physiological conditions (P7 and P30), we found no difference in mCG demethylation around *de novo* called PAM enhancers or homeostatic enhancers between tdPos vs. tdNeg microglia at P7 (**Figure 6C and 6F, Table S3**). There was a subtle difference for PAM enhancers between tdPos vs. tdNeg microglia at P30 (**Figure 6C**). However, the overall reduced fold-change of tdPos/tdNeg mCG from P7 to P30 was in the opposite direction and thus inconsistent with a model in which repression by DNA methylation regulates the microglial state switch from PAM to exPAM (**Figure 6C**). This data suggests that the DNA hypomethylation around PAM regulatory elements, presumably established during early development, may provide a permissive environment to allow rapid microglial activation later in life.

**Figure 6.**
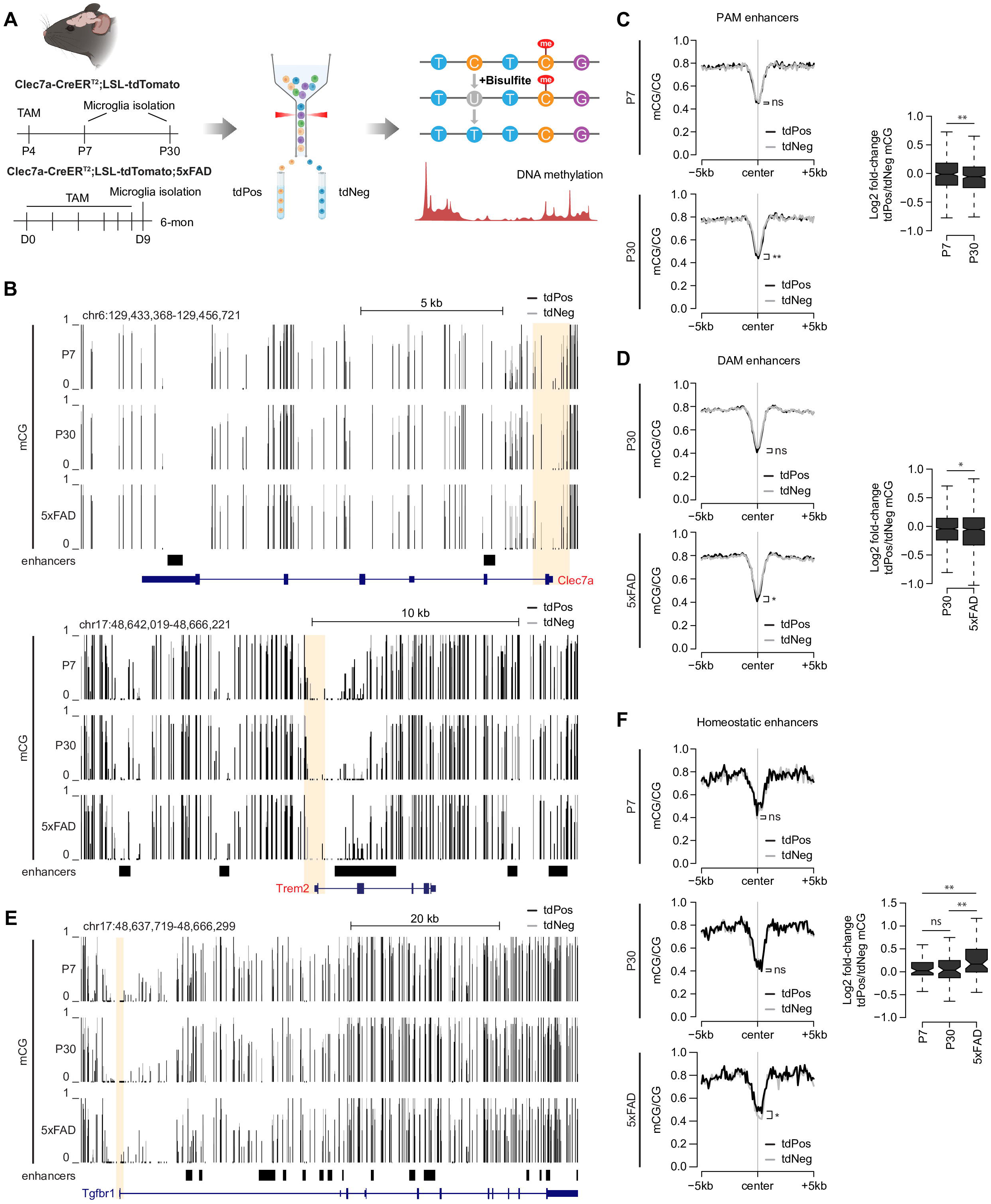
Stable DNA hypomethylation is permissive for microglial activation. **(A)** Schematic depiction of experimental design for whole genome bisulfite sequencing in P7, P30 and 5xFAD brains. TAM, tamoxifen. **(B)** Genome browser view of WGBS data for tdPos and tdNeg microglia isolated from P7 and P30 timepoints, as well as the Clec7a-CreERT2; LSL-tdTomato; 5xFAD mouse model at shared PAM and DAM upregulated DEGs, *Clec7a* and *Trem2*. Promoters are highlighted in orange. Enhancer regions are marked by black rectangular shapes. Note the hypomethylation at the promoter and enhancer regions of these genes in both tdPos and tdNeg populations from all three conditions. **(C)** Aggregate mCG levels at PAM enhancers from tdPos vs. tdNeg microglia at P7 and P30 timepoints. ** *p* < 0.01; ns, not significant; paired t-test across mean normalized CPM values between tdPos and tdNeg microglia bioreplicates (P7, n = 5; P30, n = 5). Fold-changes in mCG signals in tdPos vs. tdNeg microglia are shown on the right. ** *p* < 0.01; Wilcoxon test. **(D)** Aggregate mCG levels at DAM enhancers from tdPos vs. tdNeg microglia at P30 and in the 5xFAD model. * *p* < 0.05; ns, not significant; paired t-test across mean normalized CPM values between tdPos and tdNeg microglia bioreplicates (P30, n = 5; 5xFAD, n = 3). Fold-changes in mCG signals in tdPos vs. tdNeg microglia are shown on the right. * *p* < 0.05; Wilcoxon test. **(E)** Genome browser view of WGBS data for tdPos and tdNeg microglia isolated from P7 and P30 timepoints, as well as the Clec7a-CreERT2; LSL-tdTomato; 5xFAD mouse model at a homeostatic microglial gene, *Tgfbr1*. Promoters are highlighted in orange. Enhancer regions are marked by black rectangular shapes. Note similar hypomethylation patterns at the promoter and enhancer regions of this gene in both tdPos and tdNeg populations from all three conditions. **(F)** Aggregate mCG levels at homeostatic enhancers from tdPos vs. tdNeg microglia at P7, P30 and in the 5xFAD model. * *p* < 0.05; ns, not significant; paired t-test across mean normalized CPM values between tdPos and tdNeg microglia bioreplicates (P7, n = 5; P30, n = 5; 5xFAD, n = 3). Fold-changes in mCG signals in tdPos vs. tdNeg microglia are shown on the right. ** *p* < 0.01; ns, not significant; Wilcoxon test. **See also Figure S3, Table S3.**

In the context of AD, we observed only slightly but significantly lower mCG signals around DAM enhancers in tdPos microglia compared to tdNeg cells (**Figure 6D, Table S3**). Conversely, the tdNeg population showed slightly lower mCG signals around homeostatic enhancers as microglia transited from being in a healthy condition (P30) to disease (5xFAD) (**Figure 6F**). These changes may be attributed to longer duration of microglial activation in the AD model. However, the subtlety was in sharp contrast with the more pronounced differences observed in histone modifications at the enhancers of distinct microglial states as shown above (**Figure 4 and Figure 5**). These data suggest that while the DNA methylation pattern at enhancers is generally stable and permissive for rounds of microglial activation, it may still cooperate with histone modifications to regulate microglial states over an extensive period of time, such as during aging and progressive neurodegeneration.

### Unbiased enhancer clustering analysis reveals transcription factors that modulate PAM, DAM, and homeostatic microglial states

To obtain further biological insights into our newly identified enhancers that are associated with PAM and DAM states, we decided to systematically determine groups of co-regulated enhancers across different brain contexts in an unbiased manner. We performed a weighted gene correlation network analysis (WGCNA)^49^ using H3K27ac signal at enhancers called across all conditions (tdPos and tdNeg at P7, P30, and in 5xFAD model) (**Figure 7A, Figure S4A**). WGCNA identified 35 clusters of highly co-regulated enhancers, with 21 clusters containing at least 1000 enhancers (**Figure S4B, Table S4**). Comparison of the mean H3K27ac signal in each enhancer module across conditions revealed clusters specific to different microglial states, as well as shared enhancers (**Figure 7A**). A PAM-specific enhancer cluster ‘d’ showed the highest H3K27ac signal in tdPos microglia at P7, while DAM-specific clusters ‘i’ and ‘j’ had the highest aggregate H3K27ac signal in the tdPos population in the 5xFAD mouse model (**Figure 7A and 7B**). PAM and DAM shared enhancer clusters ‘f’ and ‘g’ had high tdPos H3K27ac signal at P7 and in the 5xFAD model compared to all other microglial populations. Finally, a homeostatic cluster ‘k’ displayed enrichment of H3K27ac signal in tdNeg microglia at P7 and in 5xFAD as well as in both P30 populations. Identification of these clusters provides defined *cis*-regulatory modules that can be explored to identify transcriptional mechanisms driving microglial states.

**Figure 7.**
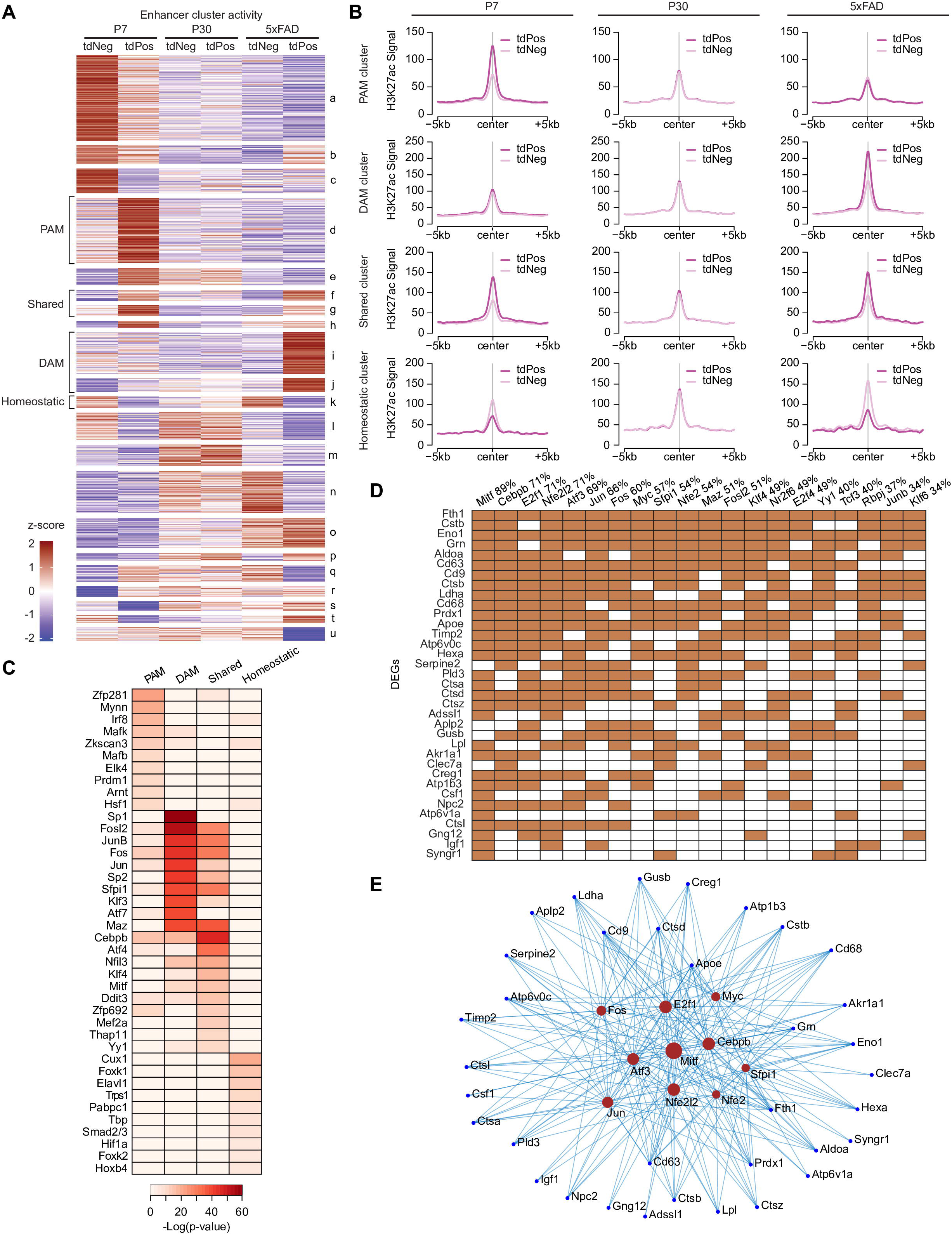
Unbiased enhancer clustering analysis leads to prediction of transcription factors regulating distinct microglial states. **(A)** Heatmap of normalized H3K27ac signals for enhancer modules identified by WGCNA across a series of conditions. **(B)** Aggregate H3K27ac levels in tdPos versus tdNeg microglia at P7, P30, and in the 5xFAD model for WGCNA enhancer clusters ‘d’ (PAM-specific), ‘i’ and ‘j’ (DAM-specific), ‘f’ and ‘g’ (shared), and ‘k’ (homeostatic). **(C)** Prediction of top 10 TFs based on enhancer-associated motifs most highly enriched in PAM, DAM, shared, or homeostatic enhancer groups by HOMER. **(D)** Prediction of interactions between TFs (based on motifs enriched in PAM and DAM shared enhancers) and PAM/DAM shared downstream DEGs by ChEA3. The numbers indicate percentage of downstream genes predicted to be regulated by a given TF candidate. **(E)** TF-target gene regulatory network highlighting potential master TF regulators of the PAM/DAM shared core gene signature. The sizes of the TFs (red) correlate with the numbers of downstream targets they regulate. **See also Figure S4, Table S4 and S5.**

Transcription factors (TFs) are critical regulators of gene expression that bind enhancer DNA elements to modulate dynamic microglial phenotypes. To infer TFs that may regulate microglial states, we performed *de novo* motif enrichment analysis using HOMER^50^ on the key enhancer clusters mentioned above. This led to the identification of sets of candidate TFs responsible for generating PAM, DAM and homeostatic microglial states (**Figure 7C, Table S5**). Notably, several top hits have been shown to play a role in regulating microglial states. For example, the *Irf8* motif was enriched in PAM-specific enhancers, and this TF has been recently reported to direct microglial transcriptional programs in the postnatal brain.^51^ *Mitf* required for DAM formation, and its motif was identified in PAM and DAM shared enhancers.^52^ Canonical TGFβ signal transduction TFs, *Smad2/3*, which promote quiescent microglial phenotypes, were predicted based on the homeostatic enhancers.^53,54^ Beyond the identification of previously implicated TFs in microglial biology that validates our approach, we also observed enrichment of additional factors potentially driving PAM/DAM formation. This includes a group of TFs related to the integrated stress response (ISR) pathway, such as ATF family genes (e.g. *Atf4*), *Ddit3* and *Nfe2l2*, which interact with one another to regulate redox and metabolic processes (**Table S5**).^55,56^ Interestingly, ISR has been implicated in AD pathogenesis, and ATF4, among other ISR genes, are elevated in AD samples.^57^ Furthermore, PU.1 (*Sfpi1*), a genetic risk factor for AD,^58,59^ was also predicted to bind and regulate PAM/DAM enhancers. Identification of these candidate drivers of reactive microglial states suggest the involvement of previously unexplored gene modules and pathways in microglia during brain development and neurodegeneration, which warrants future functional investigation.

To determine how these TFs may regulate downstream target genes, we utilized the ChEA3 (ChIP-X Enrichment Analysis Version 3) software, which is a TF enrichment analysis tool incorporating ChIP-seq and gene co-expression datasets from multiple sources, to link the candidate TFs to the DEGs from reactive microglia.^60^ We focused on TFs predicted to interact with PAM and DAM shared enhancers. This analysis demonstrated that *Mitf*, *Cebpb*, *E2f1*, *Nfe2l2* were among the most active TFs, each of which regulated over 70% of downstream target genes (**Figure 7D**). TF-target gene regulatory network also placed these genes as potential master regulators (**Figure 7E**). Taken together, these findings revealed potential *cis*- and *trans*- regulators underlying microglial state transitions in development and disease.

## DISCUSSION

Transcriptomic studies have led to the identification of numerous microglial subpopulations in physiological and pathological conditions.^15,16^ These concerted efforts leveraging unbiased *in vivo* characterization of microglial transcriptomes have refuted the oversimplified M1/M2 dichotomy of microglial reactivity and at the same time caused a nomenclature conundrum.^15,61^ On one hand, single cell profiling data are inherently noisy, making it difficult to interpret the functional implications of nuanced gene expression differences. On the other hand, each study tends to focus on one specific biological setting, leaving it open for two possibilities: (1) these different microglial states represent discrete populations of microglia generated by distinct origins and mechanisms; (2) at least some of them may be differential manifestations of reactivity by the same group of cells in response to related cues at different timepoints in life. This latter possibility insinuates *in vivo* plasticity of reactive microglial states, which has not been systematically examined. Here, we use a series of genetic fate mapping models to directly track the changes of an early postnatal microglial subpopulation, PAM, into disease and aging conditions. We demonstrate that PAM transit to a homeostatic state in adulthood, which then further convert to DAM in white matter injury and WAM in aging. Therefore, these microglial states are connected along a continuous lineage, underscoring their extraordinary plasticity.

Most tissue macrophages including microglia are thought to be derived from yolk sac precursors.^62,63^ Whether rare populations of microglia, such as PAM (less than 5% of total microglia), share the same embryonic origin is unclear. The transient appearance of PAM during the early postnatal window as well as their amoeboid morphology and the distinct transcriptional signature seem to implicate the possibility of perinatal immune infiltration.^10,30^ However, through *Runx1* lineage tracing experiment, we definitively show that PAM, just as other homeostatic microglia, are generated by the same pool of erythro-myeloid progenitors in the yolk sac. This result also indicates that PAM represent a reactive state that is triggered by developmentally regulated signals. Although we do not know the nature of these signals yet, they might be related to CNS myelination and/or ventricles due to the overlapped timing and restricted regional distribution.^10^ Future work should address this important question.

A fundamental key question in the microglial field has been to what extent reactive microglial states are plastic and reversible.^15^ Previously, we have demonstrated that DAM can fully return to homeostasis after an acute white matter injury.^17^ In this study, we focus on PAM from the developing brain to illustrate their plasticity over large timescales and across multiple conditions. Although our *Clec7a* fate mapping model would track the entire lineage of PAM cells, the BrdU/EdU co-labeling experiments demonstrate that PAM-exPAM-DAM conversions largely occur to the exact same cells rather than proliferated cells from the lineage. Interestingly, compared to non-PAM-derived cells, the PAM-derived ones seem to have no preference in the generation of DAM, suggesting a lack of immune priming in the paradigm tested. This is perhaps not surprising given that the developing brain is remarkably malleable. As inflammatory stimuli or vaccinations can produce immune memory to microglial cells in general,^18,64,65^ future work should examine the plasticity of other reactive microglial states, for instance, in a diseased brain with repetitive injuries or progressive pathologies.

An interesting observation we made about exPAM is their regional distribution. After the early postnatal development, PAM cells return to homeostasis and some of them appear to migrate out of the corpus callosum. These exPAM can be reproducibly found in superficial layers of cortex and several subcortical regions. This is consistent with the notion that perinatal white matter microglia serve as the “fountain of microglia”,^66^ except that they only contribute to a small percentage of total microglia in selective brain regions. It is unclear whether exPAM passively move along certain anatomical structures to reach these regions or they are being actively recruited there to perform some critical functions. This deserves future investigation.

In addition to characterize the plasticity of microglial states at the morphological and transcriptomic levels, we also performed efficient epigenomic profiling via CUT&Tag and WGBS to identify the enhancer landscapes and transcription factors that regulate convergent and distinct pathways across specific microglial states in development and disease. This is achieved through the utilization of a recently developed inducible driver for PAM and DAM, Clec7a-CreER^T2^, which allows acutely isolation of these rare microglial populations.^17^ Coupling this epigenomic profiling with genetic fate mapping, we find that PAM and DAM enhancers exhibit dynamic activity, reaching peak activation at P7 and in the 5xFAD model, respectively, while decreasing in activity during periods of microglial homeostasis. Interestingly, such dynamic changes of enhancer landscapes are mainly manifested in histone modifications but less so in DNA methylation. For example, all the state-associated enhancers tested are notably depleted for DNA methylation, suggesting upstream regulatory mechanisms that license these enhancers for activation later in life or in response to brain disruptions. Because histone modifications occur in the timescale of minutes to hours, which is consistent with rapid changes of microglial reactivity, whereas DNA methylation is much more stable and alterations of it tend to require extensive periods of time,^47,48,67^ it will be interesting to examine whether DNA methylation plays in role in regulating microglial states in the context of aging.

Lastly, our study uncovers key enhancer regions with distinct and shared epigenomic features that modulate PAM, DAM, and homeostatic states. This resource is particularly informative for DAM enhancers in the AD model, as many disease risk genes have been linked to microglia-specific gene and enhancer regions of the genome.^68,69^ It will be important to functionally test these enhancers to conclusively determine their contribution to AD pathogenesis. Our unbiased WGCNA supports the *de novo* differential enhancer activity analysis described above, and meanwhile it provides higher statistical power, with a majority of clusters harboring at least 1000 enhancer regions. This allows us to conduct transcription factor motif enrichment analysis to predict critical microglial state regulators. In addition to revealing the enrichment of motifs for known transcription factors associated with microglial and immune activation such as *Mitf* and NF-kappa-B complex components (e.g. *Rela*),^52,70^ which demonstrates the robustness of our analysis, we have also inferred previously unknown transcription factors associated with PAM, DAM, and homeostatic microglia states. Future studies should disrupt these regulators to assess their functional relevance on the induction and maintenance of these microglial states.

Overall, we have elucidated the plasticity of microglial states at the transcriptomic, morphological and epigenetic levels, across development, disease and aging conditions. We have identified candidate gene regulatory elements that regulate PAM in early postnatal development and DAM in neurodegenerative disease, opening up new opportunities for future functional studies. Understanding the formation and regulation of specific microglial states will shed light on fundamental mechanisms of microglial cells in shaping brain structures and modulating disease.

### Limitations of the study

Our work of characterizing microglial plasticity primarily focuses on PAM, a developmentally regulated microglial state, which may not represent other reactive states from microenvironments less prone to recovery such as neurodegeneration. Therefore, we would caution for simple generalization of our findings to other conditions. Through regional distributions of PAM and exPAM, we infer potential migration of exPAM after the early postnatal development. More direct imaging-based evidence will be needed to substantiate this conclusion. In addition, despite that the Clec7a-CreER^T2^ driver is rather specific to reactive microglial states in the brain parenchyma, its labeling efficiency is certainly not 100%. This means that in our epigenomic profiling studies, the tdTomato-negative populations may have included some *Clec7a*+ microglia that are not labeled. However, given *Clec7a*+ microglia only contribute about 3-20% of total microglia depending on the contexts, we estimate that at least 90% of tdTomato- negative microglia are *bona fide Clec7a*- microglia. This caveat might explain the level changes of histone modifications (rather than all or none) in PAM or DAM enhancers even for some unique marker genes. Related to this, for a given condition, we separate microglial states into *Clec7a*+ (reactive) vs. *Clec7a*- (homeostatic) populations. This oversimplification is justifiable due to the specificity of the Clec7a-CreER^T2^ driver and the dominant presence of homeostatic microglia in a mouse brain. However, we understand microglial reactivity is much more nuanced, and therefore, profiling of histone modification and DNA methylation at the single cell resolution will be necessary to dissect the regulatory elements underlying the full spectrum of microglial reactivity.

## ACKNOWLEGEMENTS

We thank the Alvin J. Siteman Cancer Center at Washington University School of Medicine and Barnes-Jewish Hospital in St. Louis, MO. and the Institute of Clinical and Translational Sciences (ICTS) at Washington University in St. Louis, for the use of the Genome Technology Access Center, which provided sequencing service. The Siteman Cancer Center is supported in part by an NCI Cancer Center Support Grant #P30 CA091842 and the ICTS is funded by the National Institutes of Health’s NCATS Clinical and Translational Science Award (CTSA) program grant #UL1 TR002345. The tissue clearing and imaging experiments were performed in part through the use of Washington University Center for Cellular Imaging (WUCCI) supported by Washington University School of Medicine, The Children’s Discovery Institute of Washington University and St. Louis Children’s Hospital (CDI-CORE-2015-505 and CDI-CORE- 2019-813) and the Foundation for Barnes-Jewish Hospital (3770 and 4642). Lightsheet data was generated on a Zeiss Lattice Lightsheet 7 Microscope which was purchased through support from the Arnold and Mabel Beckman Foundation as part of the Center for Advanced Lightsheet Microscopy Data Science. We thank Dr. Hiroshi Kiyonari from RIKEN Center for Biosystems Dynamics Research and Dr. Igor M. Samokhvalov for providing the Runx-Mer-Cre-Mer mouse (Accession No. CDB0524K). We thank United Kingdom Research and Innovation, specifically the MRC Harwell Archive for providing the Cdh5-CreER^T2^ mouse. We thank members from the labs of Dr. Qingyun Li and Dr. Harrison Gabel for technical support and critical reading of the manuscript. We also thank Dr. Lu Sun, Dr. Ye Zhang, and Dr. Hui Zong for helpful discussion on the project. N.H. is supported by F30HD110156. N.A. is supported by American Heart Association (#23PRE1025832). K.M.B. is supported by Ruth L. Kirschstein National Research Service Award (1F31NS139508). G.Y. is supported by the NIH (U19NS123719). H.W.G. is supported by NIH (R01NS041021 and R01AG078512). Q.L. is supported by the Whitehall Foundation (2021-08-003), BIG Center at WashU, ICTS, the Hope Center for Neurological Disorders, the McDonnell Center for Cellular and Molecular Neurobiology and NIH (R01AG078512).

## CONTACT FOR REAGENT AND RESOURCE SHARING

Further information and requests for resources and reagents should be directed to and will be fulfilled by the Lead Contact, Qingyun Li.

## AUTHOR CONTRIBUTIONS

Conceptualization, N.H., D.K., N.A., H.W.G., Q.L.; Methodology, N.H., D.K., N.A., Z.C., L.M., P.B., H.W.G., Q.L.; Software, N.H., Z.C., L.M., H.W.G., Q.L.; Investigation, N.H., D.K., N.A., Z.C., L.M., D.S., K.B., J.Y., H.W.G., Q.L.; Formal Analysis, N.H., D.K., N.A., Z.C., L.M., D.S., H.W.G., Q.L.; Writing – Original Draft, N.H., D.K., Z.C., L.M., H.W.G., Q.L.; Writing – Review & Editing, N.H., D.K., H.W.G., Q.L.; Funding Acquisition, G.Y., H.W.G., Q.L.; Resources, L.M., Z.S., P.B.; Supervision, P.B., G.Y., H.W.G., Q.L.

## DECLARATION OF INTERESTS

The authors declare no competing interest.

**See also Table S2.**

**See also Table S2.**

## STAR METHODS

### Key Resources table

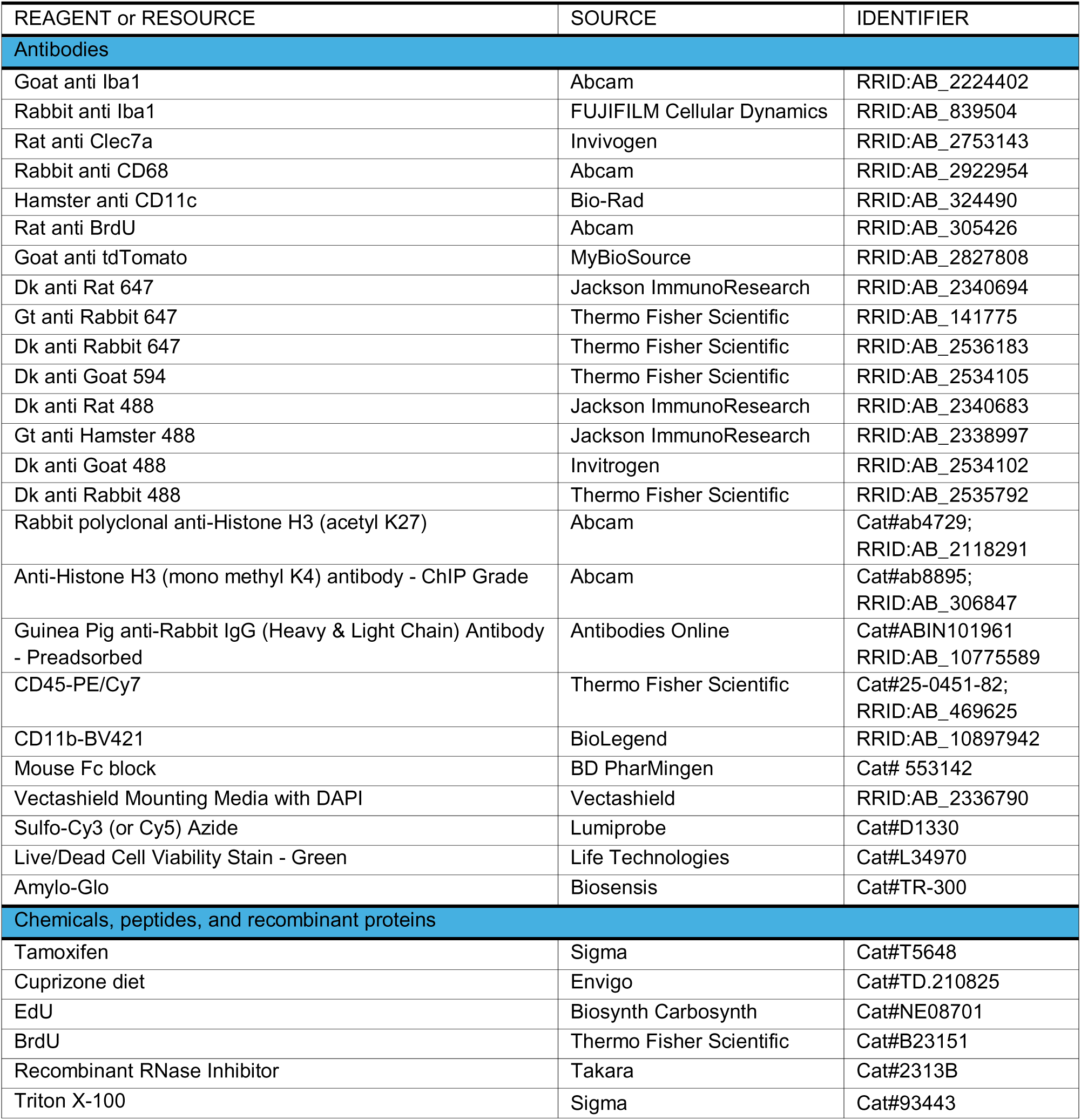
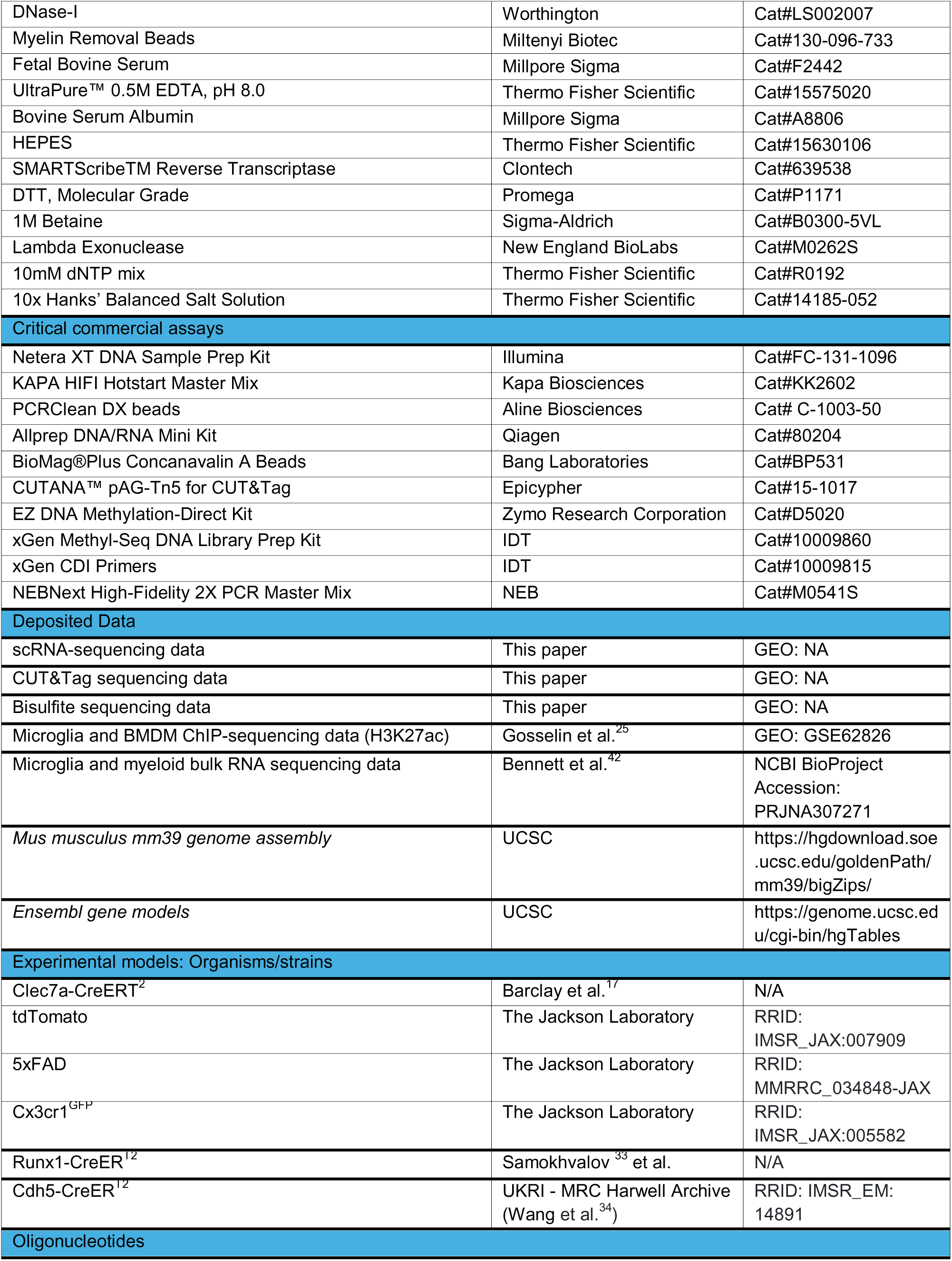
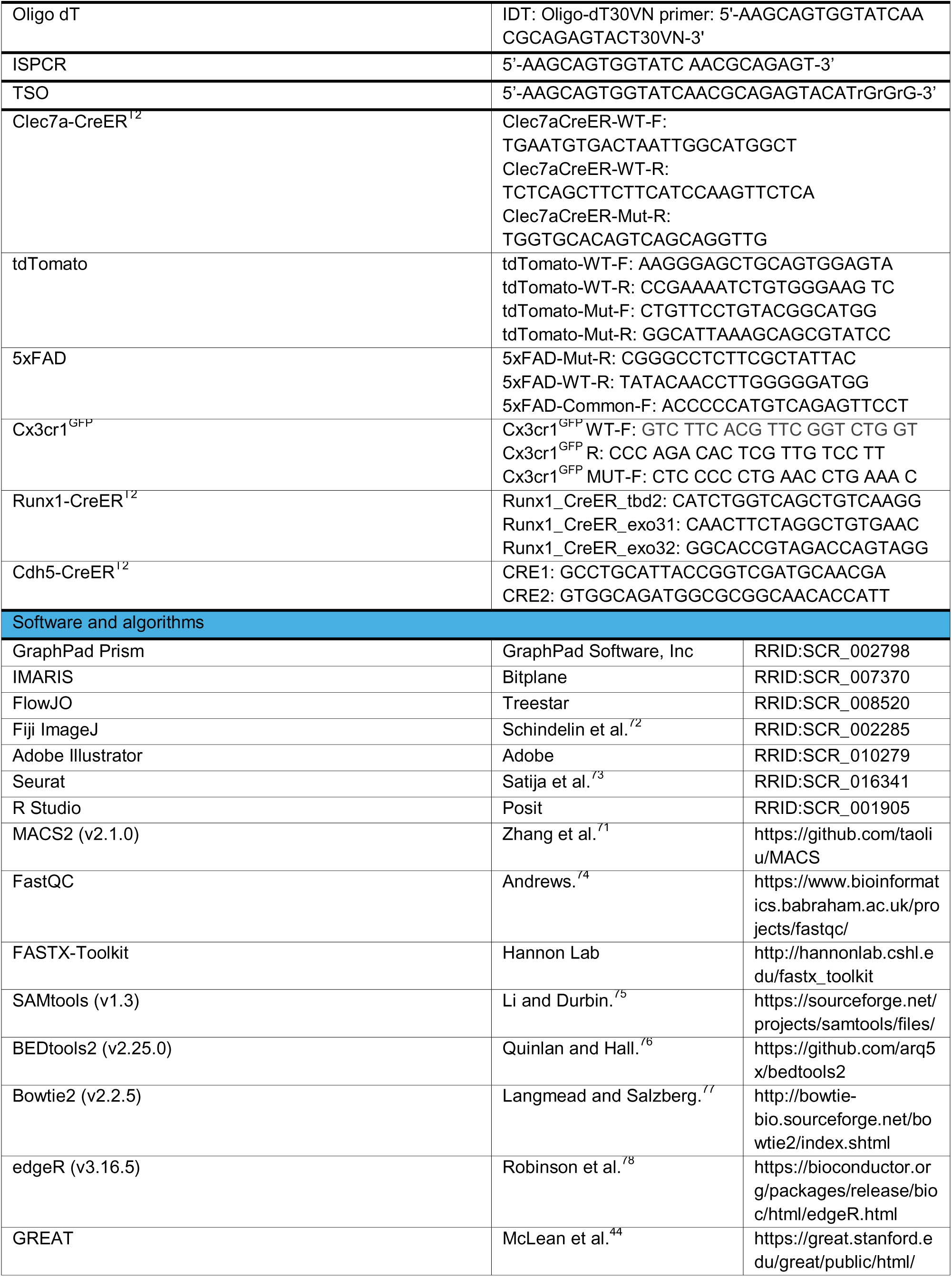
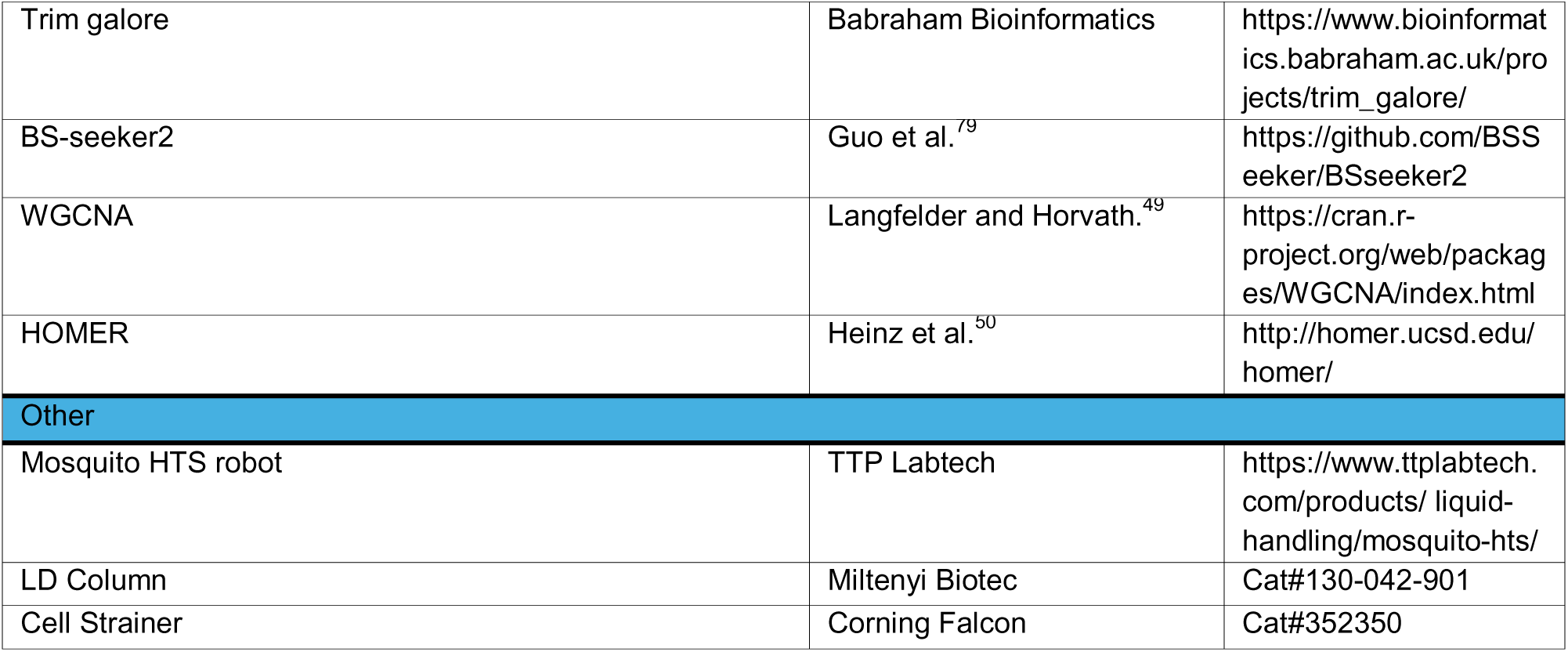

### Resource availability

#### Materials Availability

The Clec7a-P2A-iCreER^T2^ mouse is available under a material transfer agreement.

#### Data and code availability

- Single-cell RNA-seq data from P7 and P30 brains will be deposited at GEO. H3K27ac, H3K4me1, and bisulfite data for P7, P30 and 5xFAD mice will also be deposited at GEO. They will be publicly available from the date of publication. Accession numbers will be listed in the key resources table.
- This paper does not report original code.
- Any additional information required to reanalyze the data reported in this paper is available from the lead contact upon request.

### Experimental model and study participant details

#### Mice

##### Mouse lines

C57BL/6J were used as wildtype mice. Cx3cr1^GFP^ (stock #005582-JAX) and Rosa-LSL-tdTomato (Ai14) (stock #007914-JAX) mice were purchased from Jackson Laboratories. B6.Cg-Tg(APPSwFlLon,PSEN1*M146L*L286V)6799Vas/Mmjax was obtained from the Mutant Mouse Resource and Research Center (MMRRC) at The Jackson Laboratory, an NIH-funded strain repository, and was donated to the MMRRC by Robert Vassar, Ph.D., Northwestern University (RRID:MMRRC_034848-JAX).^40^ Runx1-CreER^T2^ mice were obtained from Dr. Hiroshi Kiyonari at RIKEN Center for Biosystems Dynamics Research and Dr. Igor M. Samokhvalov as the originator (Accession No. CDB0524K).^33^ Cdh5-CreER^T2^ mice were obtained from UKRI - MRC Harwell Archive (RRID: IMSR_EM: 14891).^34^ Clec7a-CreER^T2^ mice were generated in-house using CRISPR-Cas9 knock-in.^17^ All experiments were performed under the approval of the Institutional Animal Care and Use Committee at Washington University in St. Louis. Mice were housed under specific pathogen-free conditions, with controlled temperature and humidity, a 12-hour light/dark cycle, and no more than five mice per cage. Rodent chow and water were provided ad libitum. Cages and bedding were changed weekly.

### Method details

#### Tamoxifen delivery

For Runx1-CreER^T2^ and Cdh5-CreER^T2^ lineage tracing experiments, a mixture of 1.5mg (Z)-4-Hydroxytamoxifen (Sigma H7904-25MG) and 0.75mg progesterone (to counteract estrogen agonist effects of tamoxifen) prepared in 500uL corn oil was intraperitoneally administered to a pregnant mouse carrying E7.0 or E6.5 embryos respectively. For PAM labeling, a single 10uL dose of 10mg/mL tamoxifen solution prepared in corn oil was subcutaneously injected into each P4 pup. For DAM (5xFAD) labeling in 6-month-old adult mice, 200uL of 20mg/mL tamoxifen solution in corn oil was intraperitoneally injected every other day for 3 days, followed by 3 daily injections and sacrifice.

#### Cuprizone model

Clec7a-CreER^T2^; LSL-tdTomato mice were injected with tamoxifen at P4 as mentioned above and aged to 2-3 months. They were then fed with 0.2% cuprizone diet (Envig TD.210825) for 5 weeks to induce demyelination, followed by 2 weeks of control diet for spontaneous remyelination. Cuprizone diet was changed every week.

#### Immunohistochemistry

Mice were euthanized with a ketamine and xylazine cocktail prepared in saline, and then transcardially perfused using PBS, followed by 4% PFA. Extracted brains were fixed in 4% PFA overnight at 4°C and cryopreserved in 30% sucrose for 48 hours at 4°C. Brains, embedded in optimal cutting temperature medium (Tissue-Tek) were manually sectioned (50um) on a Leica cryostat.

Free-floating sagittal and coronal sections were stored in 6-well plates, blocked, and permeabilized in PBS with 10% serum and 0.2% Triton X-100 for 15 minutes at room temperature. Sections were incubated in primary antibodies diluted in PBS with 1% serum and 0.2% Triton X-100 overnight at 4°C. After three washes, sections were incubated on a shaker in secondary antibodies diluted in PBS with 1% serum and 0.2% Triton X-100 for 2 hours. Sections were washed three times, transferred onto glass slides, and mounted in Vectashield with DAPI (Vector Laboratories H-1200). For sections stained with Amylo-Glo, tissue was incubated in a 1:500 solution of Amylo-Glo diluted in PBS for 20 minutes, washed three times, transferred onto glass slides, and mounted with Vectashield (Vector Laboratories H-1000). Images were acquired using a Nikon A1R confocal microscope and quantified using Fiji software.

The following primary antibodies were used: Goat polyclonal anti-IBA1 (Abcam ab5076, 1:500); 1:1000); Goat polyclonal anti-IBA1 (FUJIFILM 019-19741, 1:500); Goat polyclonal anti-TdTomato (MyBioSource MBS448092, 1:500); Rat neutralizing monoclonal anti-mDECTIN-1-IgG (clone R1-4E4) (Invivogen mabg-mdect, 1:30); Hamster monoclonal anti-Mouse-CD11c-IgG (close N418) (Bio-Rad MCA1369, 1:40); Rabbit polyclonal anti-CD68-IgG (Abcam ab125212, 1:100); Rat monoclonal anti-BrdU (Abcam ab6326, 1:1000).

The following secondary antibodies were used: Donkey anti-goat IgG (H+L) cross-adsorbed, Alexa Fluor 488 (ThermoFisher Scientific 11055, 1:1000); Donkey anti-rabbit IgG (H+L) highly cross-adsorbed, Alexa Fluor 488 (ThermoFisher Scientific A-21206, 1:1000); Goat Anti-Armenian Hamster IgG (H+L), AffiniPure, Alexa Fluor 488 (Jackson ImmunoResearch Laboratories, 1:1000); Donkey Anti-Goat IgG (H+L) cross-adsorbed, Alexa Fluor 594 (ThermoFisher Scientific, 1:200); Donkey anti-rabbit IgG (H+L) highly cross-adsorbed, Alexa Fluor 647 (ThermoFisher Scientific 31573, 1:200); Goat anti-Rabbit IgG (H+L) highly cross-adsorbed, Alexa Fluor 647 (ThermoFisher Scientific 21245, 1:200); Donkey anti-rat IgG (H+L) AffiniPure, Alexa Fluor 647 (Jackson Immuno Research Laboratories 712-605-153, 1:200).

#### EdU (5-ethynyl-2’-deoxyuridine) labeling

EdU powder, dissolved in 100uL of 1xPBS by agitating and heating at 37°C for 30 minutes, was intraperitoneally administered at a 50mg/kg dose every three days from postnatal day 14 to day 30. Standard immunohistochemistry procedure was performed before EdU staining. Sections were then permeabilized in 0.5% PBST for 30 mins, and then incubated in an EdU development cocktail for 30 minutes at room temperature. After washing three times in 0.2% PBST, sections were incubated in 0.05% DAPI diluted in 1xPBS for 5 mins. EdU development cocktail was prepared according to the following concentrations and order listed, no more than 15 minutes before incubation: 75.75% 1XTBS (pH=7.6), 4% 100mM copper sulfate in water, 0.25% 2mM sulfo-Cy5 azide in DMSO, and 20% 500mM sodium ascorbate in water.

#### BrdU (5-bromo-2-deoxyuridine) labeling

BrdU powder (ThermoFischer Scientific, B23151), dissolved in drinking water at 0.8mg/mL was provided to Clec7a-CreER^T2^; LSL-tdTomato mice (tamoxifen injected at P4) with a concurrent 0.2% cuprizone diet for 5 weeks. Drinking water was changed every three days. Heat-mediated antigen retrieval was performed on slide-mounted samples, with 10% citrate buffer in double distilled water for 20 minutes at boiling. Slides cooled for 20 minutes before standard immunohistochemistry procedures were performed using Rat monoclonal anti-BrdU (Abcam ab6326, 1:1000).

#### Confocal image analysis

To quantify labeling efficiency in Cre reporter lines, manual counting of CLEC7A+ (or IBA1+) and tdTomato+ cells and quantification of double-positive cells were performed. To quantify labeling efficiency for EdU and BrdU samples, manual counting of CLEC7A+ and CD68+ (respectively) and tdTomato+ cells overlapping with EdU+ or BrdU+ cells and quantifying triple-positive cells were performed. Microglia morphology was quantified via manual counting of primary branches and terminal branch points. Data represents data from at least three mice. All quantifications were performed in Fiji, using the Cell Counter plugin. 3D reconstructions of microglia were generated with the Imaris software.

#### Whole-brain tissue clearing

Following perfusion, whole brains were post-fixed in 4% PFA at 4C° overnight and were washed in 1xPBS (15 mins, three times). Samples were then infiltrated with SHIELD polyepoxy^36^ for 6 days with constant agitation at 4C°. They were then polymerized at 37C° for 24 hours and were delipidated using a SmartBatch+ ETC active tissue clearing platform (LifeCanvasTechnologies, Cambridge, MA). Samples were washed in 1xPBS for 24 hours, and were refractive index matched to 1.52 using EasyIndex RI matching media (LifeCanvasTechnologies). Whole sample images were acquired on a Miltenyi Ultra Microscope Blaze gaussian lightsheet platform using a 4x/0.35 MI PLAN objective. Datasets were stitched together using Stitchy (Translucence Biosystems, Irvine, CA) and were visualized using Bitplane Imaris 10.1.1 (Oxford Instruments, Abingdon, Oxfordshire).

#### Coregistration and quantification for cleared whole brains

3D Brain coregistration onto the Allen Mouse Brain Atlas V3 was performed with the assistance of the free coregistration platform ABBA [https://abba-documentation.readthedocs.io/en/latest/].^80^ A full, detailed in-depth protocol for 3D coregistration and analysis can be found at this protocol.^81^ Briefly, 3D lightsheet brain images were first downsampled by 4x in their X and Y dimensions to enable faster processing. The brain images were then resliced perpendicularly in Fiji so that the coronal axis was now positioned along the Z-axis. The resulting coronal axis was further resampled by 10 in Python, by sampling every tenth optical slice (as opposed to interpolating), resulting in an image with 200-300 slices in the coronal axis. Both the resampled tdTomato and GFP channels were imported into ABBA, where they were first coregistered coronally in the Z-axis by matching fiducial structures, such as the hippocampus, between the lightsheet brain and the Allen Brain Atlas. Slices containing such fiducial structure were paired between image and atlas as Z-axis-immutable keyframes, matching several throughout the entire brain from the olfactory bulb through the cerebellum. A registration in Z was interpolated between these keyframes, before further coregistering each slice in the X/Y dimensions via a user-guided affine transformation^82^ that rotated, translated, and stretched the slice overtop the atlas until an accurate coregistration was obtained.

The resulting coregistered brain was exported with a region-matched atlas overlay into Fiji, during which the signal was resampled to have a pixel size of 0.02 mm per pixel in the X/Y plane, and saved as a two-channel, coregistered volume. The tdTomato channel of this export was segmented for positive-cell signal using the machine-learning aided segmentation Fiji plugin labkit.^83^ The coregistered atlas overlay was then imported atop the binarized, cell-segmented volume, where a Fiji macro was used to extract an estimate of cell-count per region, total cell-size per region, using the Fiji’s default ‘analyze particles’ function, in addition to recording region sizes. Through a Python script, this information was further used to obtain cell density per region and a clustering factor. This clustering factor represented the cellular density of any particular region divided by the cellular density of the entire brain. For example, if cells in the hippocampus were found to be five times as dense as cells throughout the entire brain, the hippocampus would have a clustering factor of 5, which in turn would denote a larger than expected proportion of cells in the hippocampus. This factor can be used to easily evaluate relative locations of cells in the brain and how they change with respect to some variable, such as developmental stage, while being robust to differing resolutions, signal intensities, and segmentation quality between samples, since the global clustering factor for a brain is normalized to a value of 1.0 between all samples.

Overall, five P7 and three P30 samples were coregistered and had clustering factors obtained for their tdTomato channels, in which PAM (or PAM-derived) cells were labelled. The clustering factors across all given regions were then individually evaluated for significant differences between the P7 and P30 cohorts, using a Wilcoxon rank-sums test, a statistical test offering high-power compared to t-test when sample sizes are low and resultingly the assumptions of the t-test cannot be met. This was performed in Python with the stats module. Before any sets of regions were compared, the clustering factors were first converted to their respective logarithms, due to the fact that the clustering factor is a multiplicative distribution but statistical testing presumes an additive distribution. Since many values had a clustering factor of 0 due to not containing segmented signal, all regions also had the value of 0.1 added to them during significance testing prior to taking their logarithm. After obtaining a final list of P7 and P30 significantly greater regions, the regions were sorted by their absolute differences in magnitudes between the average clustering factors of the P7 and P30 datapoints included for final significance testing, with the top ten for each being displayed in Figure 2B. Renders were generated in Imaris 10.1 by first creating a smoothed surface overtop a binarized Allen Mouse Brain Atlas Version 3, before importing representative segmented P7 and P30 datasets overtop the atlas render.

### Microglia isolation for FACS from brain tissue

Microglia cells were isolated using a previously published protocol.^84^ Briefly, mice were euthanized with ketamine and transcradially perfused with 20mL of 1XPBS. Choroid plexus and meninges were removed. Remaining brain tissue was finely chopped using a blade and further homogenized using a douncer and piston. Homogenized tissue solution was filtered using a 70um cell strainer, centrifuged (500g for 5 minutes at 4C°), and resuspended in MACS buffer (0.5% BSA, 2 mM EDTA in 1xPBS). Tissue solution was incubated in myelin removal beads (Miltenyi Biotec,130-096-733) and filtered through columns (LD columns for P30 and 5xFAD - Miltenyi Biotec, 130-042-901 and LS columns for P7 mice - Miltenyi Biotec, 130-042-401). The single cell suspension was centrifuged and resuspended in 1xPBS for LIVE/DEAD staining for 10 minutes (1:1000, Life Technologies, L34970). Cells were then resuspended in FACS buffer, incubated in Fc block for 5 minutes (1:60, BD Pharmingen, 553142), and primary antibodies for 10 minutes. Antibodies were washed, cells were resuspended in FACS buffer, and kept on ice for sorting. Sorting was performed using the SONY sorter (SH800) (for scRNA-seq) and Beckman CytoFLEX SRT Cell Sorter (for epigenomic profiling). Microglia were classified as CD45^int^CD11b^+^ cells, gated on live singlets. Single tdTomato+ or tdTomato- cells were sorted into 384-well plates (each well containing 0.5uL lysis buffer). Bulk tdTomato+ or tdTomato- cells were sorted into FACS tubes with FACS buffer (for CUT&Tag) or into 1mL Eppendorf tubes with RLT buffer (for bisulfite sequencing).

### scRNA-seq library preparation

Sorted microglia plates were prepared for scRNA-seq following the previously published Smart-seq2 protocol.^84^ Briefly, plates were thawed and primer annealing was performed at 72°C for 3 minutes. Reverse transcription was performed by adding 0.6uL of reverse transcription mixture (9.5U SMARTScribe Reverse Transcriptase (100U/uL, Clontech 639538), 1U RNase inhibitor (40U/uL), 1XFirst-Strand buffer, 5mM DTT, 1M Betaine, 6mM MgCl2, 1uM TSO (Rnase free HPLC purified) to each well (PCR protocol - at 42°C for 90 min, followed by 70°C, 5 min). DNA preamplification was done by adding 1.5 uL of PCR amplification mix (1XKAPA HIFI Hotstart Master Mix (Kapa Biosciences KK2602), 0.1uM ISPCR Oligo (AAGCAGTGGTATCAACGCAGAGT), 0.056U Lambda Exonuclease (5U/uL, New England BioLabs M0262S) to each well (PCR protocol - (1) 37°C 30 min; (2)95°C 3 min; (3) 23 cycles of 98°C 20 s, 67°C 15 s, 72°C 4 min; (4) 72°C 5 min). cDNA was diluted 1:10, and 0.4uL from each well was used for tagmentation.

Nextera XT DNA Sample Prep Kit (Illumina FC-131-1096) was used at 1/10 of recommendation volume, with the help of a Mosquito HTS robot for liquid transfer. Specifically, tagmentation was done in 1.6uL (1.2uL Tagment enzyme mix, 0.4uL cDNA) at 55°C, 10 min. To stop the reaction, neutralization buffer was added 0.4uL per well and incubated at room temperature for 5 min. Then 0.8uL Illumina 10-bp dual indexes (0.4uL each, 5uM) and 1.2uL PCR master mix were added to amplify whole transcriptomes using the following program: (1) 72°C 3 min; (2) 95°C 30 s; (3) 10 cycles of 95°C 10 s, 55°C 30 s, 72°C 1 min; (4) 72°C 5 min. Libraries from a single 384 plate were pooled together in an Eppendorf tube and purified twice with PCRClean DX beads. The quality and concentrations of the final mixed libraries were measured with Bioanalyzer and Qubit, respectively, before Illumina Nova sequencing.

### scRNA-seq data processing and clustering

The alignment of scRNA-seq raw data was conducted using the same pipeline as previously described.^10^ Specifically, Prinseq^85^ was utilized to filter sequencing reads shorter than 30 bp (-min_len 30), trim the first 10 bp at the 5_′_ end (-trim_left 10) of the reads, trim reads with low quality from the 3_′_ end (-trim_qual_right 25), and remove low complexity reads (-lc_method entropy, −lc_threshold 65). Afterward, Trim Galore was deployed to remove the Nextera adapters (–stringency 1), followed by STAR^86^ to align the remaining reads to the mm10 genome using the settings: – outFilterType BySJout,–outFilterMultimapNmax 20,–alignSJoverhangMin 8,– alignSJDBoverhangMin 1,–outFilterMismatchNmax 999,–outFilterMismatchNoverLmax 0.04,–alignIntronMin 20,–alignIntronMax 1000000,–alignMatesGapMax 1000000,– outSAMstrandField intronMotif. Then Picard was employed to remove the duplicate reads (VALIDATION_STRINGENCY = LENIENT, REMOVE_DUPLICATES = true). The aligned reads were converted into counts for each gene by using HTSeq (-m intersection-nonempty, -s no).^87^

The Seurat package was utilized to perform unsupervised clustering analysis on scRNA-seq data.^73^ Briefly, gene counts for cells that passed QC thresholds (percent.ribo < 0.1, percent.mt < 5, total_counts > 200,000, total_counts < 1e+7 and number_of_expressed_genes > 500, number_of_expressed_genes < 7500) were normalized to the total expression and log-transformed, and highly variable genes were detected. With the top 2000 highly variable genes as input, principal component analysis (PCA) was conducted on the scaled data, and the top components were utilized to compute the distance metric. This distance metric then underwent unsupervised cell-clustering analysis. The results were visualized in a low-dimension projection using Uniform Manifold Approximation and Projection (UMAP).^88^

### Similarity score calculation

Similarity score for scRNA-seq data was calculated as previously described.^17^ TySim^89,90^ was utilized to quantify to what extent each individual cell is similar to a specific target cell type (e.g., PAM or homeostatic microglia). Briefly, scRNA-seq data was separated by TySim into binary part and non-zero part, which were handled separately so as to mitigate the impact of drop-out effects in scRNA-seq datasets. In each part of data, artifact factors were taken into account by TySim. These factors could jointly influence observed expression values in scRNA-seq data, such as variations in sequencing depth across cells and heterogeneous preferences for different genes during sequencing. The background expression attributable to these artifacts for each gene within each cell was also considered. These background expression levels were precisely estimated by systematically modeling both cell and gene factors embedded within the input scRNA-seq data, a process achieved by employing the Conditional Multifactorial Contingency (CMC) model. In this study, the signatures of PAM and homeostatic microglia were obtained from previously published papers.^17^ The code of TySim can be obtained from https://github.com/yu-lab-vt/CMC/tree/CMC-TySim.

### Cleavage under targets and tagmentation (CUT&Tag) from sorted microglia

FACS purified tdNeg or tdPos microglia were centrifuged for 5 min at 600xg at 4°C. Cells were resuspended in 100uL/sample cold Nuclear Extraction (NE) Buffer (20mM HEPES-KOH, pH 7.9; 10mM KCl; 0.5mM Spermidine; 0.1% Triton-X 100 20% Glycerol; 1x protease inhibitor cocktail) and incubated on ice for 10 min. BioMagPlus Concanavalin A coated beads (Bangs Laboratories BP531) were prepared by washing 5uL/sample beads with Bead Activation Buffer (20mM HEPES, pH 7.9; 10mM KCl; 1mM CaCl2; 1mM MnCl2) twice, and subsequently resuspending in 5uL/sample Bead Activation Buffer and incubated at RT for 15 min. 100uL of nuclei/sample were pipetted into PCR-strip tube containing 5uL activated beads and incubated at RT for 10 min. Unbound supernatant was removed, and bead-bound nuclei were resuspended in 50uL Antibody Buffer (Wash150 Buffer (20mM HEPES, pH 7.5; 150mM NaCl; 0.5mM Spermidine; 1x protease inhibitor cocktail); 2mM EDTA) with 0.01ug of appropriate primary antibody (H3K27ac ab4729, H3K4me1 ab8895) and incubated on a nutator overnight at 4°C. Supernatant was removed, and bead-bound nuclei resuspended in 50uL cold DIG150 Buffer (Wash150 Buffer; 0.01% Digitonin) with 0.5ug secondary antibody (Guinea Pig anti-Rabbit IgG ABIN101961) and incubate at RT for 30 min on nutator. Supernatant was removed, and bead-bound nuclei washed once with cold 200uL DIG150 Buffer. Bead-bound nuclei was resuspended in 50uL cold DIG300 Buffer (Wash300 Buffer (20mM HEPES, pH 7.5; 300mM NaCl; 0.5mM Spermidine; 1x protease inhibitor cocktail); 0.01% Digitonin) with 0.5ug CUTANA pAg-Tn5 enzyme and incubated at RT for 1 hr on nutator. Supernatant was removed, and bead-bound nuclei was washed once with 200uL cold DIG300 Buffer. Supernatant was removed, and bead-bound nuclei was resuspended in 50uL Tagmentation Buffer (Wash300 Buffer; 10mM MgCl2) and incubated at 37°C for 1 hr. Supernatant was removed, and bead-bound nuclei resuspended in 50uL TAPS Buffer (10mM TAPS, pH 8.5; 0.2mM EDTA). Supernatant was removed, and 5uL of RT SDS Release Buffer (10mM TAPS, pH 8.5; 0.1% SDS) was added to bead-bound nuclei. Samples were vortexed for 7-10sec and incubated for 1 hr at 58°C. 15uL RT SDS Quench Buffer (0.67% Triton-X 100 in molecular grade H_2_O) was added to each sample. 2uL of appropriate indexing primers added to each sample. 25uL NEBNext High-Fidelity 2X PCR Master Mix (cat#M0541S) was added to each sample. CUT&Tag libraries were amplified under the following PCR cycling parameters: 58°C, 5 min; 72°C, 5 min; 98°C, 45 sec; 98°C, 15 sec; 60°C, 10 sec; 72°C, 1 min; 4°C hold. Libraries were PCR-amplified for 18-23 cycles depending on starting microglia cell input. Libraries were purified using 1.3 volumes of AmpureXP Beads. Libraries were then pooled to a final concentration of 5nM and sequenced using the Illumina Novaseq 6000 with the Genome Technology Access Center at Washington University in St. Louis, targeting 5-10 million paired-end reads per sample.

### Whole genome bisulfite sequencing (WGBS)

DNA was isolated from 10,000 FACS purified tdNeg or tdPos microglia per condition using the Allprep DNA/RNA Mini Kit (QIAGEN). Libraries were prepared for sequencing using the IDT xGen Methyl-Seq DNA Library Prep Kit (cat# 10009860) and the EZ DNA Methylation-Direct Kit (Zymo, D5020) was used for bisulfite conversion. 4-10 ng of DNA was fragmented for 45 sec with the Covaris S220 sonicator (10% Duty Factory, 175 Peak Incidence Power, 200 cycles per burst, microTUBE 200µL AFA Fiber). DNA was then purified using 0.7 volumes of SPRISelect Beads (Beckman Coulter Life Sciences) to select for long DNA inserts for sequencing. Samples underwent bisulfite conversion under the following cycling conditions: 98°C, 8 min; 64°C, 4.25 h; 4°C hold. Libraries were PCR-amplified for 15 cycles. Libraries were then pooled to a final concentration of 5nM and sequenced using the Illumina Novaseq 6000 with the Genome Technology Access Center at Washington University in St. Louis, targeting 100 million paired-end reads per sample.

### Identification of enhancers

Enhancers in this study were defined by the presence of H3K4me1 peaks that occur outside of a known promoter region. Promoter regions were defined as a 1kb region surrounding the TSS (+/-500bp). This led to the inclusion of some subthreshold regions of H3K27ac enrichment that may not represent true active regulatory elements, but ensured that we analyzed enhancers across all regulatory states (inactive, primed, active) in our studies. For this analysis, bed files of H3K4me1 CUT&Tag were pooled by replicate and across all conditions/timepoints (P7, P30, 5xFAD) for tdNeg and tdPos microglia separately. Peaks of H3K4me1 CUT&Tag were identified using the MACS2 peak calling algorithm on the pooled bed files using macs2 callpeak --keep-dup all -q 0.05. tdNeg and tdPos peak files were then combined and overlapping peaks were merged into single peaks using bedtools merge. Bedtools intersect was used to identify H3K4me1 peaks that did not overlap with gene promoter regions (1kb around annotated TSS). H3K4me1 peaks that remained were defined as enhancers.

### CUT&Tag analysis

Sequenced reads were trimmed with fastx_trimmer −l 75 and mapped to mm39 using bowtie2 alignment parameters --local --very-sensitive-local --no-unal --no-mixed --no-discordant --phred33 -I 10 -X 700. Reads were deduplicated and extended to 250bp size based on average library read length sizes. For paired-end sequenced libraries, BED files for reads 1 and 2 were concatenated prior to CUT&Tag signal quantification. Bedtools coverage –counts was used to quantify CUT&Tag signal at genomic regions. edgeR on H3K4me1 (Log2FC >0, FDR <0.05) and H3K27ac (Log2FC >0) CUT&Tag signals was used to determine differentially active enhancer signals between tdNeg and tdPos groups across conditions/timepoints.

### WGBS analysis

For bisulfite sequencing analysis, data were adaptor-trimmed using Trim Galore, and subsequently aligned to mm39, deduplicated, and called for methylation using BS- seeker2 (with bowtie2). Methylation levels across regions were assessed using bedtools map -o sum, summing the number of reads mapping to Cs (supporting methylated C, mC) and the number of reads mapping to Cs + Ts (supporting unmethylated C, C) in the region, then dividing those two numbers to calculate the absolute level or percent methylation (mC/C) in each region. Regions were adjusted for methylation nonconversion rates as measured by % methylation in Lambda spike-ins per-sample, as previously described.^91^ Following %mC calculation, the lambda % methylation value was subtracted from the calculated %mC value. If the corrected value was below 0, the %mC value was set to 0.

### Enhancer-gene linkage using GREAT

To calculate the significance and magnitude of the linkage between differentially expressed genes and differentially active enhancers, differentially expressed genes were resampled based on their expression levels, and a Fisher’s exact test was performed comparing the association of enhancers with the observed differentially expressed gene set and the resampled, expression-matched gene set. This process was repeated 1000 times, and the observed odds ratio (OR) versus the mean OR across 1000 resamplings was plotted.

### Weight gene correlation network analysis (WGCNA)

WGCNA was performed using CPM normalized H3K27ac CUT&Tag signal in tdNeg and tdPos microglial populations across all timepoints and conditions (P7, P30, 5xFAD). blockwiseModules() was ran with the following adjusted parameters: power = 10, minModuleSize = 20, maxBlockSize = 30000, #should max out for your ram (30,000), mergeCutHeight = 0.225.

### Transcription factor (TF) motif enrichment analysis using HOMER

*De novo* TF motif analysis was performed using findMotifsGenome.pl -noweight -nlen 0 -size 200 -len 8 with resampled enhancer regions based on acetylation levels as background (-bg). Motif enrichments were scored using HOMER’s default binomial distribution framework.

### TF Downstream targets prediction

To predict the downstream targets of the candidate TFs (based on PAM and DAM shared enhancers), TF–target associations were retrieved from the ChEA3 database, which integrates data from multiple sources, including TF–gene co-expression, ChIP-seq experiments, and TF–gene co-occurrence.^60^ Corresponding target genes for the driving TFs were extracted from these associations.

Recognizing that TF–target interactions are cell-type specific, the target gene list was refined to those highly expressed in both PAM and DAM cells. To identify these highly expressed genes, differential gene expression analysis was performed using scRNA-seq data and cluster labels from the previously published dataset.^17^ Specifically, clusters “DAM 1” and “DAM 2” were compared to “Adult Brain Homeostatic Microglia” to identified DEGs in DAM cells, and the “PAM” cluster was compared to “Early Postnatal Homeostatic Microglia” to obtain DEGs for PAM cells. In the analysis, the FindMarkers function in the Seurat R package was used with thresholds (p_adj < 0.05, logfc.threshold = 0.25, and min.pct = 0.1). The intersection of DEGs from both analyses yielded genes highly expressed in both cell states.

The refined TF–target associations were visualized using a heatmap that displayed relationships between the driving TFs and their target genes. In the heatmap, TFs and genes were ordered based on the number of their association.

### TF-Target genes regulatory network

To better visualize the regulatory patterns, a TF–target gene regulatory network was constructed based on the refined TF–target associations mentioned above. In this network, each node represents a TF or a gene, and nodes are connected by edges if there is an association between them. To avoid an overly complex visualization, only the top 10 TFs with the most associations were included. The node size for each TF was set proportional to the number of their associated genes to highlight potential key regulators.

### Quantification and Statistical Analysis

To quantify the fluorescence intensity of CLEC7A expression in P7, P10, P11, and P12 wild-type pups (at least 3 animals per condition), tissue sections were imaged by confocal microscopy. Fluorescence intensity from the relevant channel was measured with Fiji by subtracting background signals from area integrated intensity. To quantify microglial density, the number of cells was manually counted in Fiji, divided by areas in mm^2^. Summary statistics was calculated (*p*<0.05 cutoff) for fluorescence and density at each developmental age, one-way ANOVA followed by Tukey’s multiple comparison post hoc test was performed on GraphPad prism. Data were represented by mean +/- SEM. To calculate mean labeling efficiency and morphology, 3 IHC sections (50um) from 3 animals (10 microglial cells per animal for morphology quantifications) were analyzed using Fiji and GraphPad Prism. Statistical significance (*p* < 0.05) for each measurement was calculated by either one-way ANOVA followed by Tukey’s multiple comparison post hoc test or Student’s t-test. Summarized data were represented by mean +/- SEM. To calculate statistical significance levels for similarity score calculation from scRNA-seq data, we first computed similarity scores for each individual cell relative to the target cell type, and then performed t-test on these scores to assess statistical significance. Statistical analyses related to the epigenomic data were described in relevant figure legends.

## Supplemental Information

Document S1. Figures S1-S4.

**Table S1.** Quantification showing brain regions with the greatest differences in clustering of tdTomato+ microglia in P7 and P30, **related to Figure 2**.

**Table S2.** TdPos and tdNeg microglia edgeR CUT&Tag signal outputs at *de novo* enhancers across P7, P30, and 5xFAD conditions, **related to Figures 3-5**.

**Table S3.** TdPos and tdNeg microglia CG DNA methylation levels at *de novo* enhancers across P7, P30, and 5xFAD conditions, **related to Figure 6**

**Table S4.** Table of WGCNA cluster outputs at *de novo* enhancers, **related to Figure 7**.

**Table S5.** TF predictions based on HOMER motif analysis (most significantly enriched TFs in a given enhancer set are highlighted in red), **related to Figure 7**.

**Video S1.** Light sheet microscopy images for P7 whole brain (Clec7a-CreER^T2^; tdTomato in red, CX3CR1-GFP in green), spinning movie, **related to Figure 2**.

**Video S2.** Light sheet microscopy images for P7 whole brain (Clec7a-CreER^T2^; tdTomato in red, CX3CR1-GFP in green), sagittal optical sections, **related to Figure 2**.

**Video S3.** Light sheet microscopy images for P30 whole brain (Clec7a-CreER^T2^; tdTomato in red; CX3CR1-GFP in green), spinning movie, **related to Figure 2**.

**Video S4.** Light sheet microscopy images for P30 whole brain (Clec7a-CreER^T2^; tdTomato in red; CX3CR1-GFP in green), sagittal optical sections, **related to Figure 2**.

