## Supplemental Figures for "Microglial plasticity governed by state-specific enhancer landscapes"

Figure S1

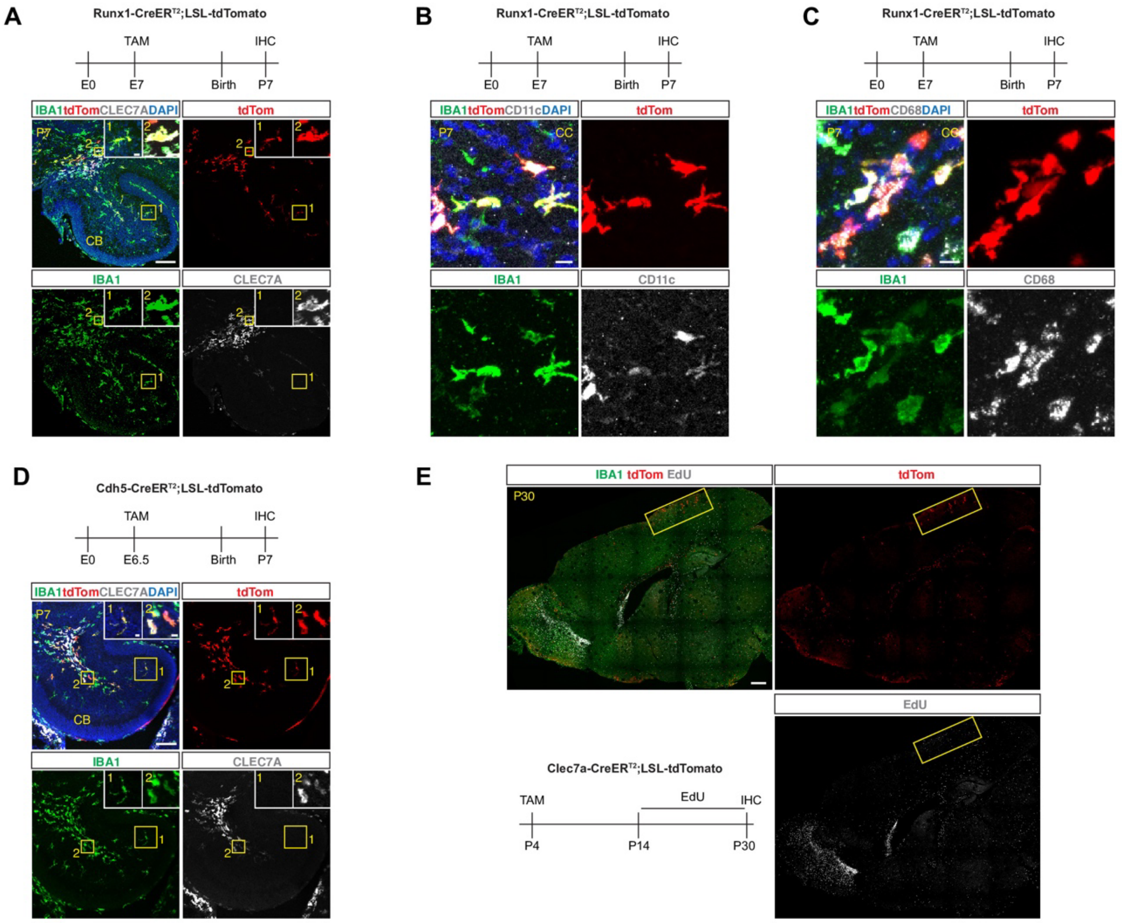


**Figure S1. Embryonic origin of PAM and their limited proliferation after the first two postnatal weeks**

**(A)** Representative immunostaining images showing tdTomato+ microglia co-stained with CLEC7A in the CB, cerebellum, of a P7 mouse brain in the Runx1-creER; LSL-Tdtomato lineage tracing model. Boxed inserts show zoomed-in images of microglia in numbered areas. Tamoxifen regimen is shown on the top. Scale bars, 100um for large images and 10um for inserts.

**(B)** Representative immunostaining images showing tdTomato+ microglia co-stained with CD11c in the CC, corpus callosum of a P7 mouse brain in the Runx1-creER; LSL-Tdtomato lineage tracing model. Tamoxifen regimen is shown on the top. Scale bars, 10um.

**(C)** Representative immunostaining images showing tdTomato+ microglia co-stained with CD68 in the CC, corpus callosum of a P7 mouse brain in the Runx1-creER; LSL-Tdtomato lineage tracing model. Tamoxifen regimen is shown on the top. Scale bars, 10um.

**(D)** Representative immunostaining images showing tdTomato+ microglia co-stained with CLEC7A in the CB, cerebellum, of a P7 mouse brain in the Cdh5-CreER$^{T2}$; LSL-tdTomato lineage tracing model. Boxed inserts showing zoomed-in images of microglia in numbered areas. Tamoxifen regimen is shown on the top. Scale bars, Scale bars, 100um for large images and 10um for inserts.

**(E)** Representative immunostaining images showing tdTomato+ microglia co-labeled with EdU in a P30 brain of Clec7a-CreER$^{T2}$; LSL-tdTomato lineage tracing mice. Note that a vast majority of EdU+ cells (e.g. along the rostral migratory stream) do not overlap with tdTomato signals. The boxed cortical region is enlarged and shown in Fig 1J. Tamoxifen regimen is shown on the bottom left.  Scale bars, 500um.

### Figure S2

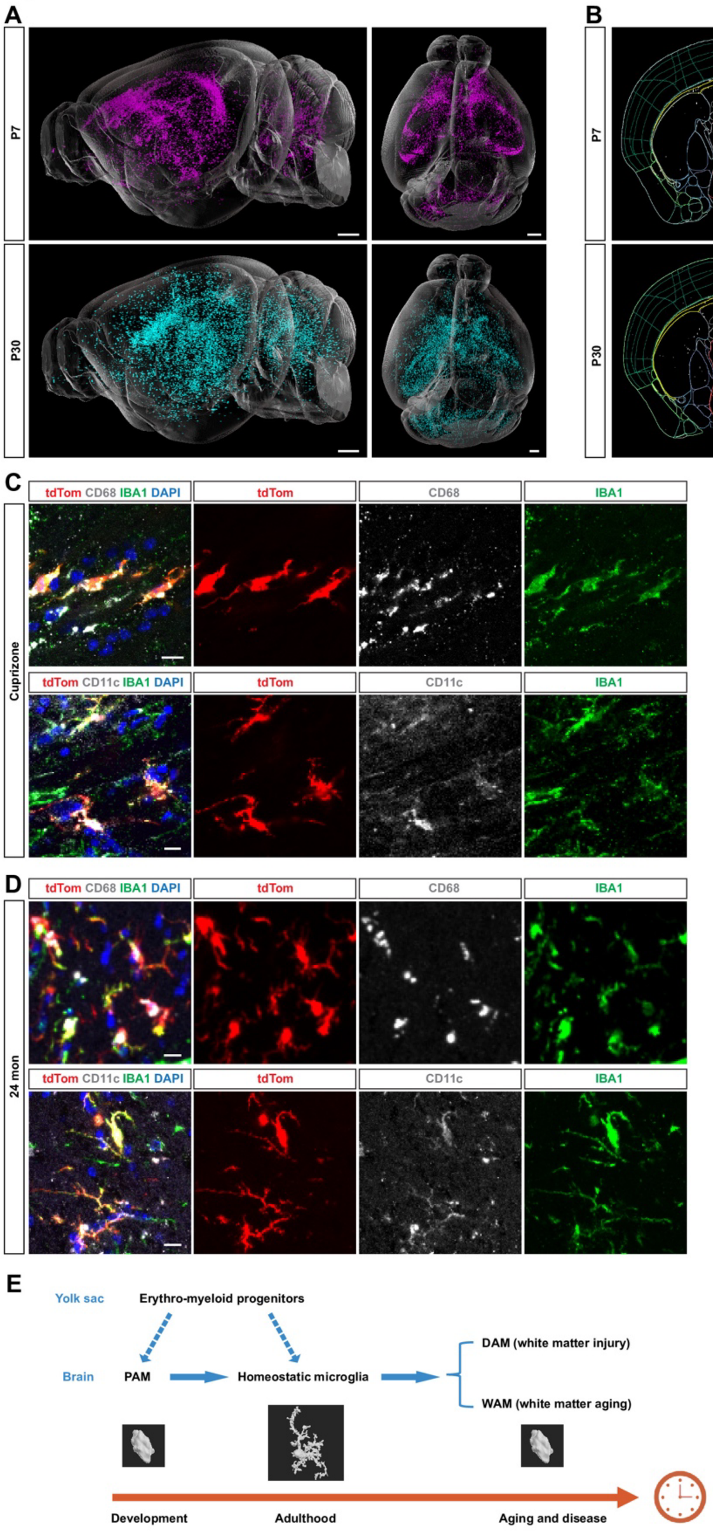

A
P7
P30
B
CC
STR
C
Cuprizone
tdTom CD68 IBA1 DAPI
tdTom
CD68
IBA1
tdTom CD11c IBA1 DAPI
CD11c
Clec7a-CreER$^{T2}$;LSL-tdTomato
Cuprizone or ctrl diet
TAM
5 wks
IHC
P4
8 wks
13 wks
D
24 mon
P4
E
Yolk sac
Erythro-myeloid progenitors
Brain
PAM
Homeostatic microglia
DAM (white matter injury)
WAM (white matter aging)
Development
Adulthood
Aging and disease

**Figure S2. Plasticity of PAM during injury and aging**

**(A)** Lateral and dorsal view of tdTomato+ microglia from cleared whole brains of Clec7a-CreER$^{T2}$; LSL-tdTomato; CX3CR1-GFP mice projected onto a standard brain model. P7 microglia are shown in magenta on the top and P30 microglia are shown in cyan on the bottom. Tamoxifen injection is done at P4. Only the tdTomato signals are shown. Scale bars, 1000um.

**(B)** Coronal optical section of cleared whole brains of Clec7a-CreER$^{T2}$; LSL-tdTomato; CX3CR1-GFP mice projected onto a standard brain atlas, showing higher enrichment of P7 tdTomato+ microglia in the CC, corpus callosum and higher enrichment of P30 tdTomato+ microglia in the STR, striatum region (CC and STR regions are outlined yellow). The arrow points to P7 tdTomato+ microglia also enriched in certain areas of layer 6, as shown by the quantification in Figure 2C. Scale bars, 1000um.

**(C)** Representative immunostaining images showing tdTomato+ microglia co-stained with CD68 (top) and CD11c (bottom) in the corpus callosum of cuprizone-treated mice in the Clec7a-CreER$^{T2}$; LSL-tdTomato model. Tamoxifen and cuprizone regimens are shown on the right. Scale bars, 10um.

**(D)** Representative immunostaining images showing tdTomato+ microglia co-stained with CD68 (top) and CD11c (bottom) in the corpus callosum of 24-month-old mice in the Clec7a-CreER$^{T2}$; LSL-tdTomato model. Tamoxifen regimen is shown on the right. Scale bars, 10um.

**(E)** Schematic graph showing that proliferative-region-associated microglia (PAM) originating from yolk sac erythro-myeloid progenitors convert to homeostatic microglia after early postnatal development and become disease-associated microglia (DAM) in response to white matter injury or white matter-associated microglia (WAM) in aging. Morphological differences of these microglial states are exemplified by 3D models.

Figure S3

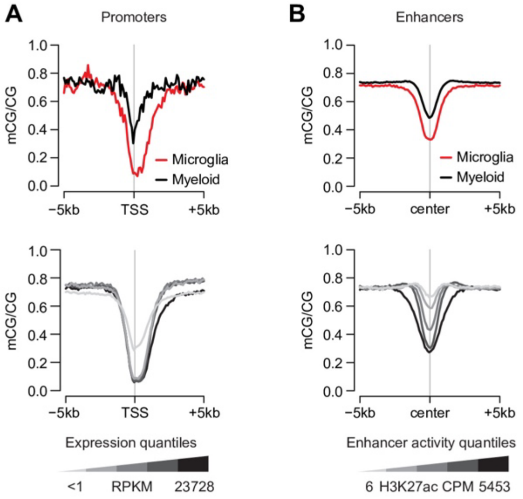


**Figure S3. WGBS captures DNA methylation levels at microglia-specific promoters and enhancers. Related to Figure 6.**

**(A)** Aggregate DNA methylation (mCG) signals at microglial gene promoters versus myeloid gene promoters (top left) and across quantiles of microglial gene expression (bottom left).

**(B)** Aggregate mCG signals at microglial enhancers versus myeloid enhancers, and across quantiles of microglial enhancer H3K27ac levels (right). Microglia and myeloid gene expression and enhancer data were from the same source as in Figure 3B.

### Figure S4

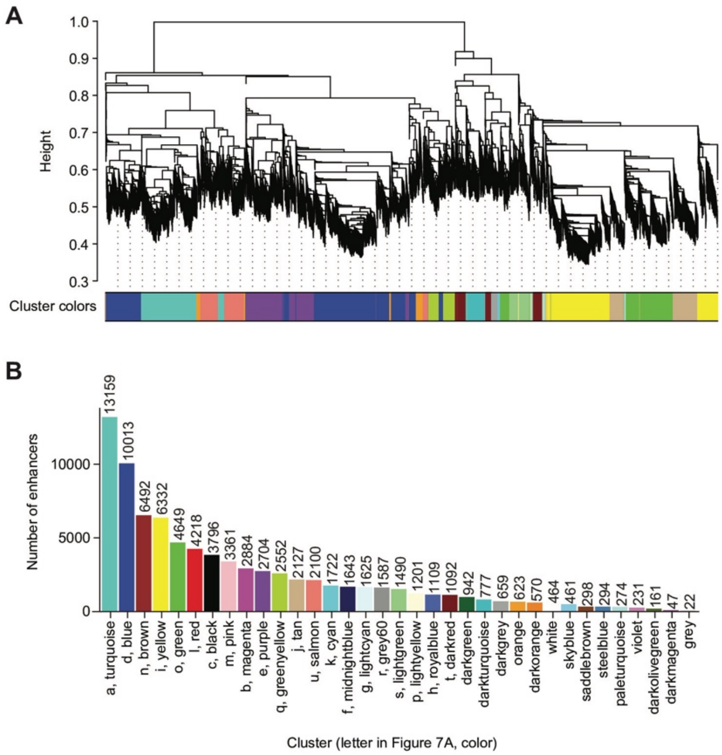


**Figure S4. WGCNA uncovers distinct microglial enhancer groups. Related to Figure 7.**

**(A)** Dendrogram of enhancers identifying WGCNA clusters (see *methods*).

**(B)** Barplot of number of enhancers in each WGCNA cluster.
